# Effects of access condition on substance use disorder-like phenotypes in male and female rats self-administering MDPV or cocaine

**DOI:** 10.1101/2024.03.04.583431

**Authors:** Michelle R. Doyle, Nina M. Beltran, Mark S. A. Bushnell, Maaz Syed, Valeria Acosta, Marisa Desai, Kenner C. Rice, Katherine M. Serafine, Georgianna G. Gould, Lynette C. Daws, Gregory T. Collins

## Abstract

Substance use disorder (SUD) is a heterogeneous disorder, where severity, symptoms, and patterns of substance use vary across individuals. Yet, when rats are allowed to self-administer drugs such as cocaine under short-access conditions, their behavior tends to be well-regulated and homogeneous in nature; though individual differences can emerge when rats are provided long– or intermittent-access to cocaine. In contrast to cocaine, significant individual differences emerge when rats are allowed to self-administer 3,4-methylenedioxypyrovalerone (MDPV), even under short-access conditions, wherein ∼30% of rats rapidly transition to high levels of drug-taking. This study assessed the SUD-like phenotypes of male and female Sprague Dawley rats self-administering MDPV (0.032 mg/kg/infusion) or cocaine (0.32 mg/kg/infusion) by comparing level of drug intake, responding during periods of signaled drug unavailability, and sensitivity to footshock punishment to test the hypotheses that: (1) under short-access conditions, rats that self-administer MDPV will exhibit a more robust SUD-like phenotype than rats that self-administered cocaine; (2) female rats will have a more severe phenotype than male rats; and (3) compared to short-access, long– and intermittent-access to MDPV or cocaine self-administration will result in a more robust SUD-like phenotype. After short-access, rats that self-administered MDPV exhibited a more severe phenotype than rats that self-administered cocaine. Though long– and intermittent-access to cocaine and MDPV self-administration altered drug-taking patterns, manipulating access conditions did not systematically alter their SUD-like phenotype. Evidence from behavioral and quantitative autoradiography studies suggest that these differences are unlikely due to changes in expression levels of dopamine transporter, dopamine D_2_ or D_3_ receptors, or 5-HT_1B_, 5-HT _2A_, or 5-HT_2C_ receptors, though these possibilities cannot be ruled out. These results show that the phenotype exhibited by rats self-administering MDPV differs from that observed for rats self-administering cocaine, and suggests that individuals that use MDPV and/or related cathinones may be at greater risk for developing a SUD, and that short-access MDPV self-administration may provide a useful method to understand the factors that mediate the transition to problematic or disordered substance use in humans.

## Introduction

The use of psychoactive substances has been commonplace for millennia, yet only 15-30% of individuals that use these drugs develop a substance use disorder (SUD) [1]. The DSM-5 defines SUD by 11 diagnostic criteria from four different modalities [2], resulting in a heterogeneous disorder, the severity of which depends on the number of positive criteria [1]. In recent decades, preclinical SUD research has attempted to model the heterogeneity and multi-symptomatic complexities of SUD in animals by assessing multiple behavioral endpoints thought to be related to the diagnostic criteria for SUD [3–6]. These multifaceted studies aim to provide greater insight into SUD and help close the translational gap in the development of effective treatments.

The 3-criteria model, developed by Deroche-Gamonet and colleagues, was one of the first attempts to quantitatively assess the SUD-like phenotypes of rats self-administering cocaine based upon (1) the number of responses made during signaled periods of drug unavailability (“drug-seeking”), (2) breakpoints under progressive ratio schedule of reinforcement (“motivation to use drug”), and (3) resistance to footshock punishment (“continued use despite adverse consequences”) [3]. Since then, this model has been adapted and used in numerous studies to identify and assess differences between rats with robust and mild SUD-like phenotypes [4–9]. Initial studies used short-access procedures, where drug was typically available during 1-2 hour sessions; however, subsequent studies suggest that providing longer periods of access to drug (e.g., 6 hrs; long-access) [10] can enhance resistance to footshock punishment [11–13], and increases in cue– and drug-induced reinstatement of extinguished responding [14–16]. More recently, an intermittent-access procedure (5-min of drug availability provided every 30-min over a 6-hour session) was developed to establish rapid, binge-like patterns of cocaine intake [17,18] and has been shown to increase the reinforcing effectiveness of cocaine, and enhance cue– and drug-induced reinstatement of responding [6,19,20]. Thus, while it is possible to observe individual differences in SUD-related behaviors in rats self-administering cocaine (or other drugs) under short-access conditions, mounting evidence suggests when rats are provided long– or intermittent-access to cocaine they develop more severe SUD-like phenotypes (for review, see [21]).

3,4-methylenedioxypyrovalerone (MDPV) is a synthetic cathinone that functions as a cocaine-like monoamine uptake inhibitor, but unlike cocaine which is roughly equipotent at the dopamine, norepinephrine, and serotonin transporters (DAT, NET, and SERT, respectively) MDPV is ∼800-fold selective for DAT and NET over SERT [22–24]. We have previously reported that 30-40% of male and female rats that are allowed to self-administer MDPV during daily 90-min sessions rapidly develop a high-responder phenotype, characterized by levels of MDPV intake ∼2-5 times greater than low-responders across a range of doses, greater breakpoints under progressive ratio schedules of reinforcement, and higher rates of responding during periods of signaled drug unavailability [25–29]. Though consistent with MDPV high-responder rats engaging in SUD-like behaviors, it is unclear if the phenotype observed in these rats would extend to other core SUD-related behaviors, such as resistance to punishment by footshock, (i.e., “continued use despite adverse consequences”), or if the “severity” of the SUD-like phenotype is sensitive to access condition manipulations, as has been reported for cocaine. Thus, the primary goals of the current study were to: (1) directly compare the SUD-like phenotype of rats self-administering MDPV to those of rats self-administering cocaine under short-access conditions; (2) determine whether long– and intermittent-access to MDPV and cocaine self-administration differentially impacted the SUD-like phenotypes relative to rats maintained on short-access MDPV and cocaine self-administration; and (3) assess whether any of these effects differed as a function of sex.

In addition, because stimulant use is known to dysregulate dopaminergic and serotonergic systems, the current studies also assessed how various drug histories (e.g., MDPV vs. cocaine, short-, long– and intermittent-access), and ultimately the severity of SUD-like phenotype impacted the expression of key transporters and receptors within the dopamine and serotonin systems. Specifically, we used quantitative autoradiography within the caudate putamen and nucleus accumbens to investigate transporters and receptors that have been shown to be increased (i.e., DAT and dopamine D_3_ receptors [30–43]; SERT and 5-HT_2A_ receptors [44–47]) or decreased (i.e., dopamine D_2_ receptors [48–56], but see [57,58]; 5-HT_1B_ and 5-HT_2C_ receptors [59–61]) following periods of stimulant use. Thus, the overarching goals of these studies were to determine the degree to which the severity of a SUD-like phenotype was influenced by the self-administered drug (MDPV and cocaine), access conditions associated with SUD-like behaviors (short-, long-, and intermittent-access), and the sex of the subject (female and male), and whether key neurobiological changes in dopamine and serotonin systems differed as a function of their SUD-like phenotype severity. Ultimately by identifying conditions that facilitate the development of robust SUD-like phenotypes, these studies will inform future work aimed at identifying medications capable of normalizing aberrant drug-taking behavior in the hopes of developing novel and effective treatments for stimulant and other substance use disorders.

## Materials and Methods

### Subjects

Female and male Sprague Dawley rats (weighing 200-225g and 275-300g, respectively, upon arrival) were obtained from Envigo (Indianapolis, IN, USA) and singly housed in a temperature– and humidity-controlled environment under a 14/10-hour light cycle (lights on at 06:00) with *ad libitum* access to Purina chow and water. All experiments were conducted in accordance with the Institutional Animal Care and Use Committee of the University of Texas Health Science Center at San Antonio, and the Guide for Care and Use of Laboratory Animals [62].

### Surgery

Rats were anesthetized using 2% isoflurane and surgically prepared with a chronic indwelling catheter in the left femoral vein, which was attached to a vascular access button secured in the mid-scapular region, as previously described [26–28,63,64]. Penicillin G (60,000 U/rat) or Excede (20 mg/kg) was administered subcutaneously following surgery, and catheters were flushed daily with 0.5 ml heparinized saline (100 U/ml) to maintain catheter patency.

## Self**-**Administration

### Apparatus

Intravenous drug self-administration was conducted in standard operant chambers (Med Associates Inc, St. Albans, VT) within light– and sound-attenuated cubicles. A white house light was located on the top of the wall opposite the two levers. Above each lever was a set of red, yellow, and green LEDs. The grid floor was connected to a scrambled shock system (Env-414, Aversive stimulator/scrambler; Med Associates Inc, St. Albans, VT) used to deliver footshocks. A variable-speed syringe driver was used to deliver infusions through Tygon tubing that was connected to a fluid swivel and spring tether held by a counterbalanced arm. The active lever (counterbalanced across rats) was signaled by illumination of the yellow LED above the lever; completion of the response requirement (fixed ratio [FR] 1 or 5) resulted in delivery of the drug infusion and initiation of the 5-sec post-infusion timeout (TO), signaled by illumination of the houselight and all three LEDs above the active lever.

### Experimental overview

As shown in Figure 1, rats initially underwent a self-administration training period where they were allowed to self-administer cocaine (0.32 mg/kg/infusion) or MDPV (0.032 mg/kg/infusion) for 24 sessions, with the first 14 sessions being under a fixed ratio (FR) 1:TO 5-sec schedule of reinforcement, and the remaining 10 being a FR5:TO 5-sec schedule. The doses were selected because of their position on the descending limb of the FR dose response curve and due to the 10-fold potency difference [22,28,29,65,66]. Then rats underwent a two-part phenotyping procedure where four endpoints were measured to generate a phenotype score. After this, rats were assigned to self-administer under short (FR5:TO 5-sec; 60-min session), long (FR5:TO 5-sec; 6-hr session), or intermittent access (FR5:TO 1.5-sec; 5 min of drug availability followed by 25-min of drug unavailability in a 6-hr session) for 3 weeks before going through the phenotyping period a second time to test the effects of access condition. Finally, rats underwent a 3-week drug-free period before doing a cue reactivity test where drug was signaled to be available, but only saline was delivered upon completion of the FR5:TO 5-sec schedule of reinforcement. Dopamine D_3_ receptor sensitivity was also measured using pramipexole-induced yawning (see Supplemental Methods) before self-administration began and at the end of the study, prior to euthanasia for quantitative autoradiography studies (see Supplemental Methods).

**Figure 1.**
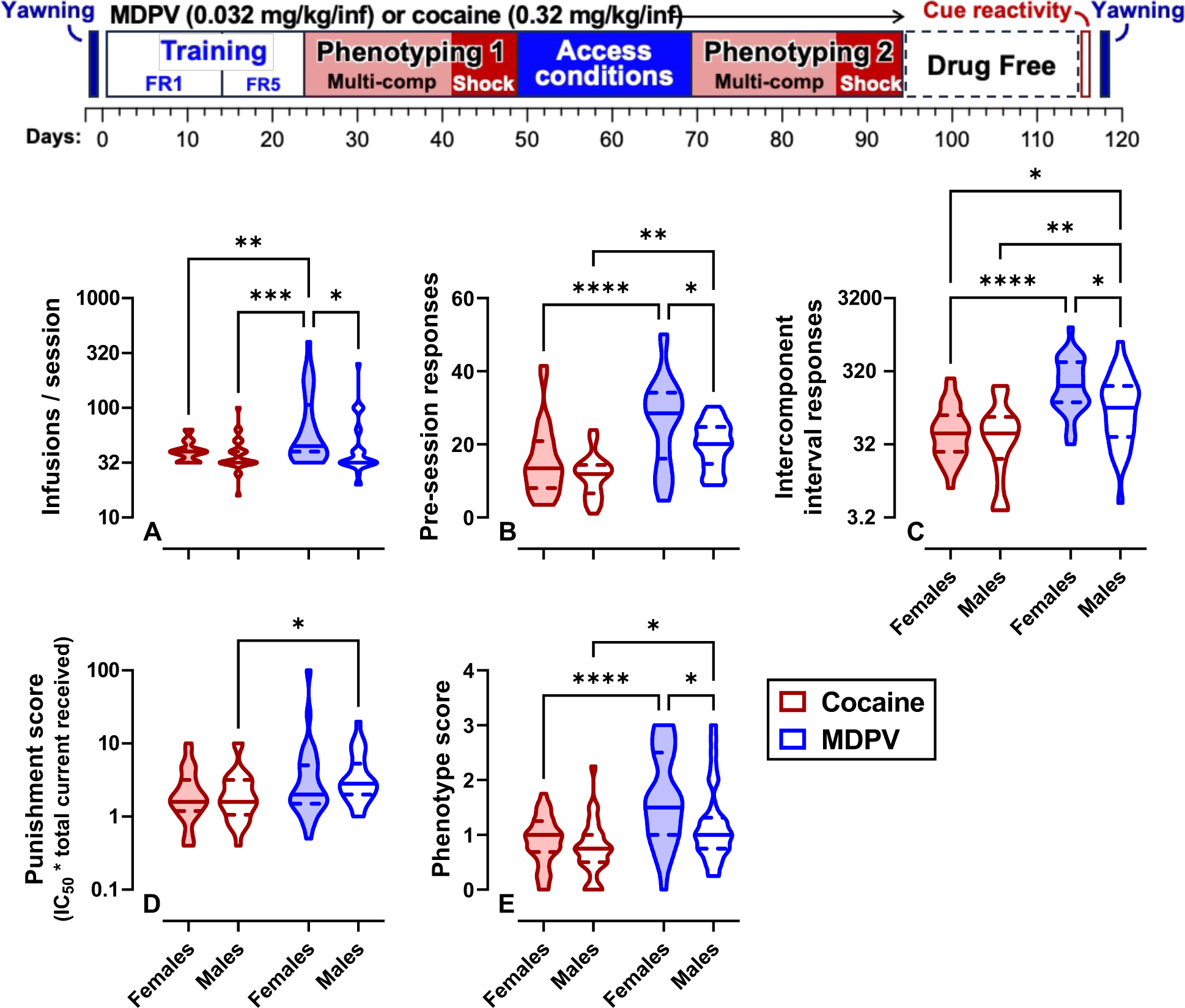
Timeline and Initial Phenotype Score Endpoints Experimental timeline showing the total study duration in days as well as each aspect of the experiment. Violin plots representing the mean number of infusions (A), pre-session responses (B), intercomponent interval responses (C), punishment score (D) and SUD-like phenotype score (E) in female (shaded) and male (white) rats self-administering cocaine (red) or MPDV (blue) during the first phenotyping period. Solid lines indicate median and dashed lines indicate quartiles. * =P<0.05, **=P<0.01, ***=P<0.001, ****P=<0.0001 for post-hoc analyses.

### Phenotyping procedure

To test the effects of access condition on an overall phenotype score, rats underwent a phenotyping procedure before and after the three-week access condition manipulation. The overall phenotype score was a composite score from four endpoints: (1) number of infusions, (2) pre-session responses, (3) intercomponent interval responses, and (4) punishment score. The first three endpoints were collected during the multiple component self-administration and the punishment score was generated during the footshock punishment procedure (see below and Figure 1 for more details).

To generate a phenotype score, all rats, regardless of sex or drug, were rank ordered for each endpoint (i.e., pre-session responses, infusions, intercomponent interval responses, and punishment score) and divided by quartiles. Rats in the bottom quartile received a score of 0, rats middle two quartiles received a score of 1, and rats in the top quartile received a score of 2; statistical outliers received scores of 3 [(1.5 x interquartile range) + 3^rd^ quartile cutoff] or 4 [(3 x interquartile range) + 3^rd^ quartile cutoff]. Individual phenotype scores represent the mean of the four endpoint scores, with the lowest possible score being 0, and highest possible score being 4. Rats with an overall phenotype score of less than 1 were classified as “low score”, rats with a score ≥1 but <2 were classified as “mid score”, and rats with a score of ≥2 were classified as having a “high score”.

### Phenotyping procedure: Multiple component self-administration

To assess responding during periods of signaled drug availability (and unavailability), all rats were transitioned to a multiple-component schedule of reinforcement that began with a 5-min pre-session TO, followed by three 20-min periods of drug availability, each followed by 5-min intercomponent TOs. During the pre-session and intercomponent TOs, drug was signaled to be unavailable by extinguishing all visual stimuli, and responses were recorded but had no scheduled consequence. A yellow LED above the active lever was used to signal drug availability. The total number of infusions earned, responses made during the presession TO, and responses during the three intercomponent TOs (i.e., periods of drug unavailability) served as endpoints for the phenotype score.

### Phenotyping procedure: Footshock punishment

To assess sensitivity to footshock punished responding, all rats responded under an FR5:TO 5-sec schedule for two 60-min “baseline” sessions before initiating footshock testing. Punishment sessions were identical to “baseline” sessions with the exception that beginning with the 4^th^ infusion, a 0.5-sec, unsignaled footshock was delivered coincident with 1 out of every 2 infusions (i.e., ∼50% of infusions were paired with a footshock). The initial footshock intensity was 0.1 mA, and this increased by 0.2 mA across consecutive sessions (0.1, 0.3, 0.5, 0.7mA) until the number of infusions earned was ≤20% of baseline, or a maximum intensity of 0.7mA (Figure 1). Punishment sessions were followed by at least two “baseline” sessions. A punishment score was calculated by multiplying the current that reduced responding by 50% (IC_50_) with the total infusions earned, and this was fourth endpoint of the phenotype score.

### Observation of behavioral response to non-contingent footshock

To determine if sensitivity to footshock differed across groups, rats were allowed to habituate to a self-administration chamber for 2-5 minutes before receiving a series of non-contingent footshocks. Behavioral responses were scored by a trained observer using the following criteria: 0=no reaction; 1=looks around or passive movement (no startle response); 2=runs around or walks backward rapidly; 3=jump; 4=vocalize. Footshock functions were generated in triplicate, twice in ascending order, and once in descending order. Scores were averaged across the three replicates.

### Drugs

Racemic MDPV was synthesized and supplied by Kenner Rice and cocaine hydrochloride was provided by National Institute on Drug Abuse Drug Supply Program. Both self-administered drugs were dissolved in sterile, physiological saline and delivered intravenously at a volume of 0.1 ml/kg and a dose of 0.032 mg/kg/infusion for MDPV and 0.32 mg/kg/infusion for cocaine.

### Statistical analyses

A one-factor (score) ANOVA was used for rate and level of acquisition with Tukey’s post-hoc analyses (Table 1). Raw data for infusions, intercomponent TO responses, punishment score, and rate of responding were log-transformed before analyzing using two-factor or three-factor ANVOAs. Two-factor or three-factor ANOVAs were performed on the raw values for pre-session responses, cue reactivity responses, phenotype score, and change in phenotype score. Two-factor (drug x sex) ANOVAs were conducted for the dataset in Figure 1, and three-factor (drug x sex x access condition or drug x sex x phenotype score) ANOVAs were performed on datasets in Figures 2-4. Tukey’s post-hoc analyses were performed when there was a significant main effect of access condition or phenotype score. Change in phenotype score (Figure 3F) was analyzed by calculating the mean change in phenotype score (phenotype score 2 – phenotype score 1) and comparing 95% confidence intervals, which were corrected for multiple comparisons. Escalation (Table 2) was initially calculated in individual subjects (mean of the last three sessions – mean of first 3 sessions of the access condition manipulation). The mean and 95% confidence intervals were used to compare whether there was significant escalation (confidence intervals did not overlap with 0). The behavioral response to noncontingent shock was analyzed using a two-factor (shock intensity x group) ANOVA, where group was phenotype score, self-administered drug, or sex. Sidak’s multiple comparisons post-hoc analyses were performed when there was a main effect of group. Data from a subset of rats (n=8; n=2 per sex/drug) were excluded from the cue reactivity test due to procedural error.

**Figure 2.**
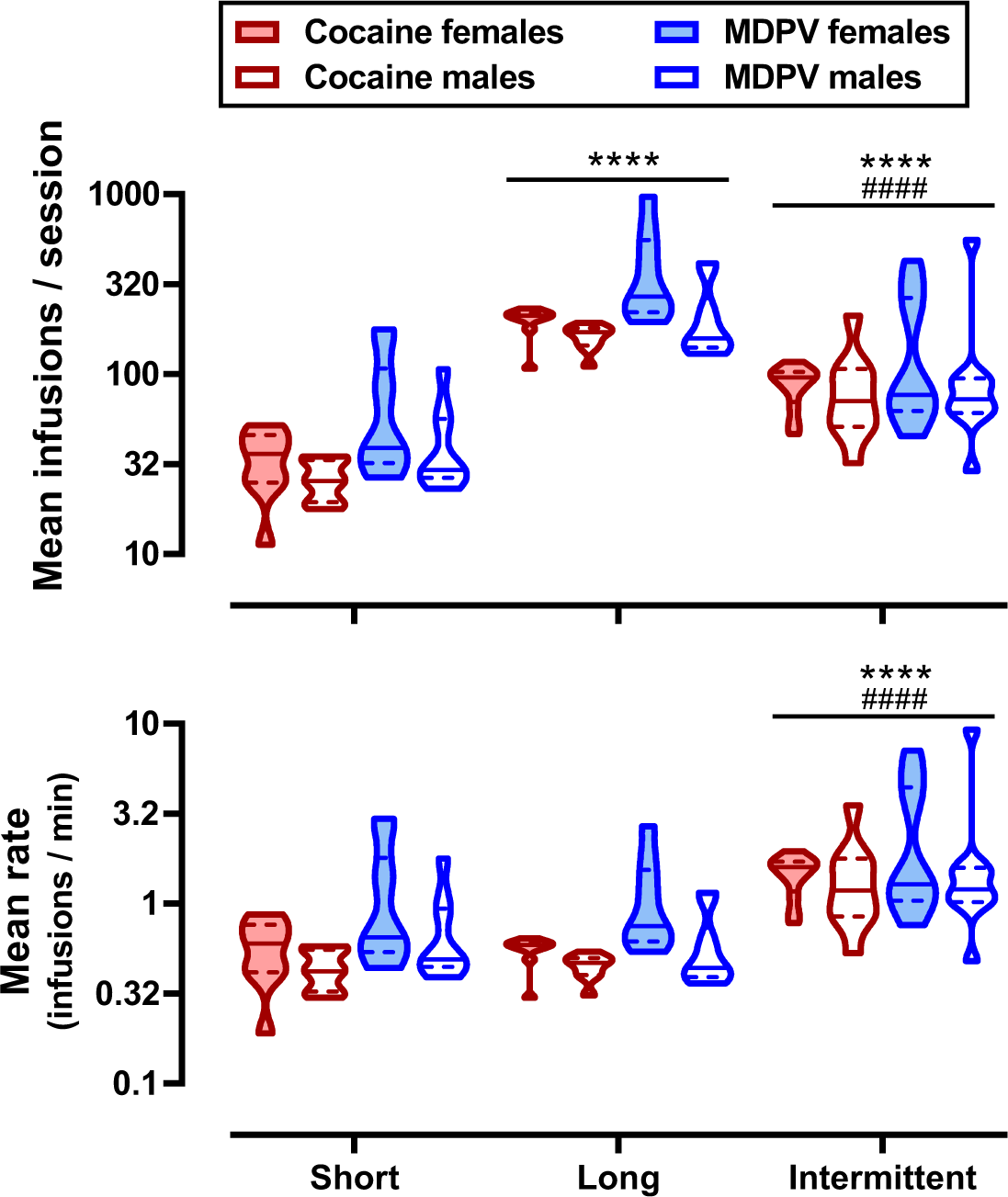
Access Condition Manipulations Violin plots representing the number of infusions (top) and rate of responding (bottom) averaged across the 21-session access condition manipulation. Female (shaded) and male (white) rats self-administering cocaine (red) or MPDV (blue) under short-(left), long-(middle), or intermittent-access (right). Solid lines indicate median and dashed lines indicate quartiles. Main effect of access condition where ****=P<0.0001 compared to short-access; ####=P<0.0001 compared to long-access for post-hoc analyses.

**Figure 3.**
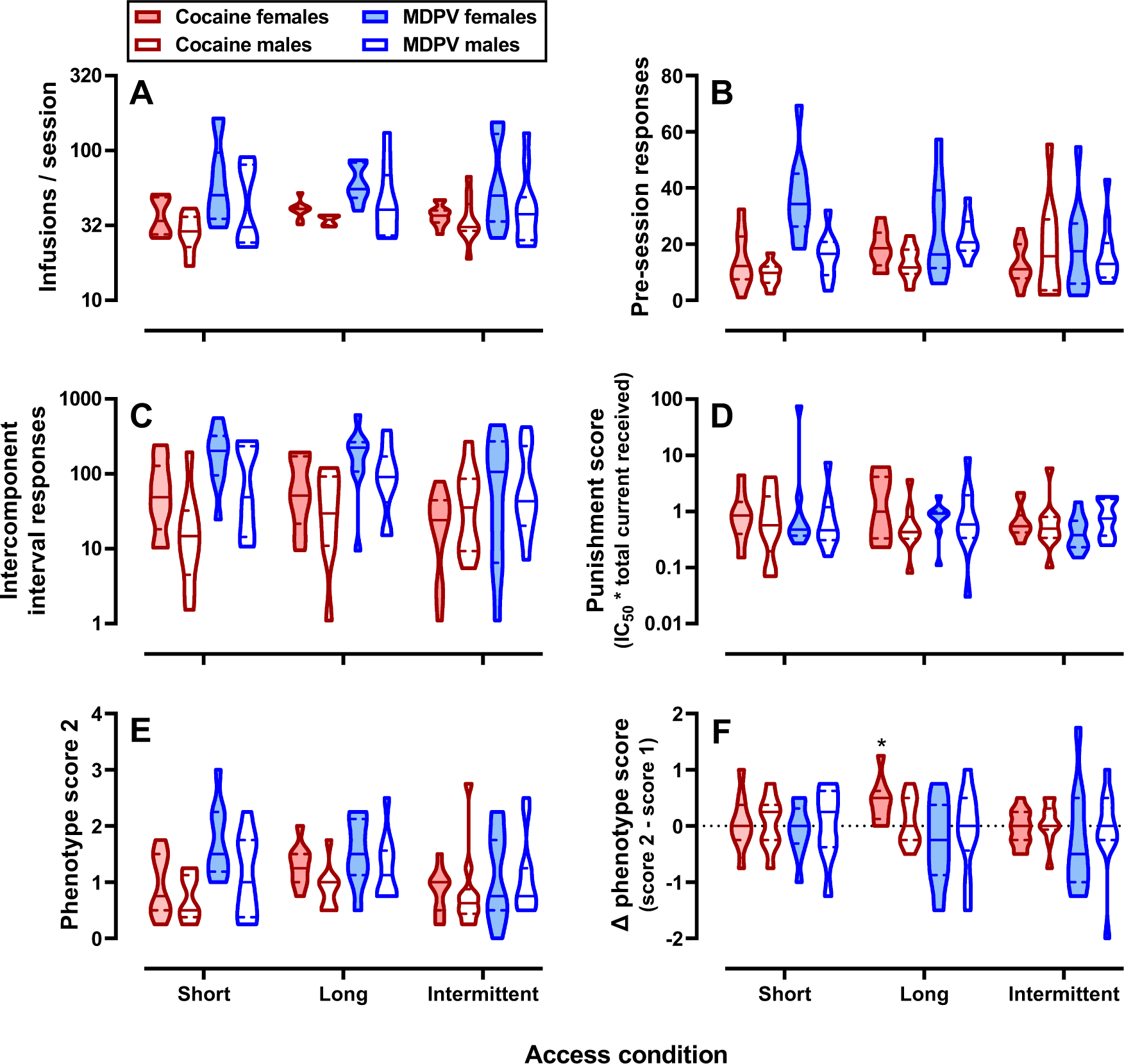
Final Phenotype Score Endpoints Violin plots representing the mean number of infusions (A), pre-session responses (B), intercomponent interval responses (C), punishment score (D), SUD-like phenotype score (E), and change in phenotype score in female (shaded) and male (white) rats self-administering cocaine (red) or MPDV (blue) during the second phenotyping period. Data are split by access condition. Solid lines indicate median and dashed lines indicate quartiles. *=significant change in phenotype score, where confidence intervals did not overlap with 0. Main effects of drug and sex are not indicated on figures.

**Figure 4.**
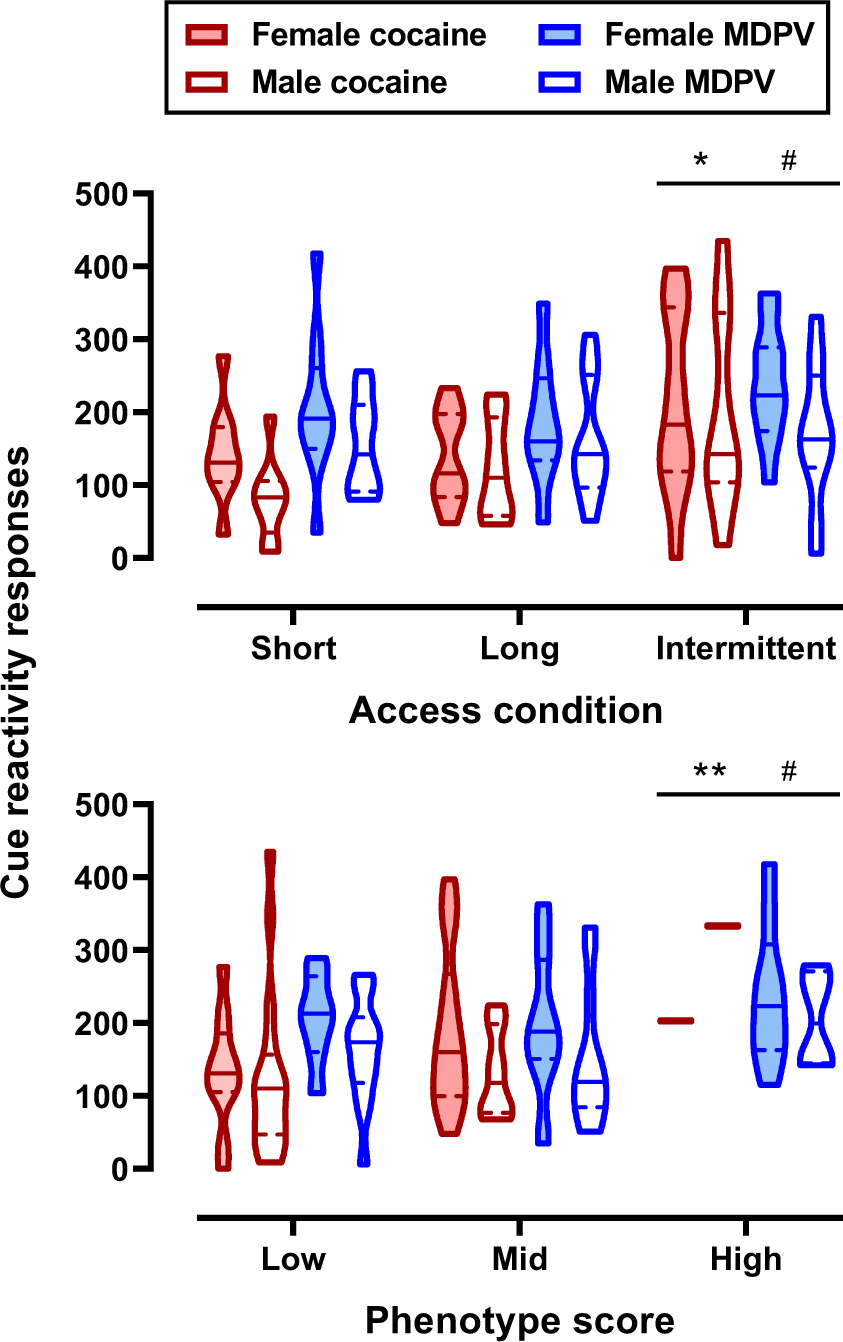
Responses During Cue Reactivity Test Violin plots representing the mean number of responses during the cue reactivity test, split by access condition (top) or phenotype score (bottom) in female (shaded) and male (white) rats self-administering cocaine (red) or MPDV (blue). Solid lines indicate median and dashed lines indicate quartiles. Top: main effect of access condition where *=P<0.05 compared to short-access; #=P<0.05 compared to long-access. Bottom: main effect of phenotype where **=P<0.01 compared to low score; #=P<0.05 compared to mid score.

**Table 1.**
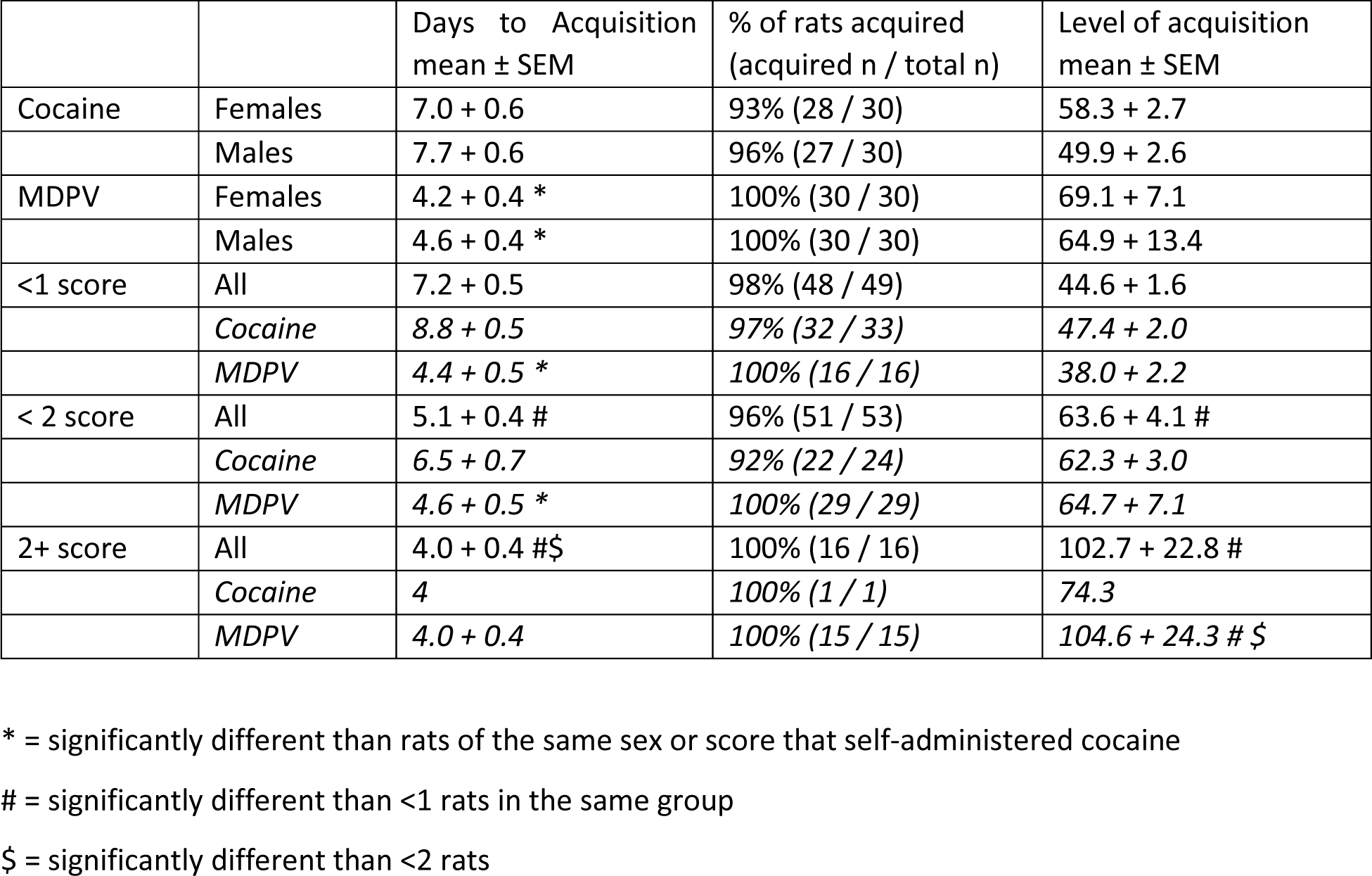
Acquisition of Cocaine and MDPV Self-Administration. The mean number of days to reach acquisition criteria (>20 infusions and >80% of responses on the active lever), percent of rats that acquired, and level of acquisition (mean infusions / session) in rats, split by sex, self-administration drug and initial phenotype score.

**Table 2.** Escalation During 21-Day Access Condition Manipulation. Mean and confidence intervals (CI) around escalation, calculated by mean of infusions earned during sessions 19-21 minus mean of sessions 1-3 in individual subjects. Data shown by self-administration group and phenotype score.

| Access condition | Group | n | Escalation<br>Mean (95% CI) |
| --- | --- | --- | --- |
| Short | Female cocaine | 9 | 0.6 (-8.5, 9.6) |
|  | Male cocaine | 9 | 0.8 (-2.8, 4.4) |
|  | Female MDPV | 10 | -11.3 (-50.0, 27.4) |
|  | Male MDPV | 9 | 1.1 (-25.3, 27.5) |
| Long | Female cocaine | 9 | **39.8 (21.3, 58.2) \*** |
|  | Male cocaine | 9 | **45.0 (12.7, 77.3) \*** |
|  | Female MDPV | 9 | 160.8 (-141.1, 462.7) |
|  | Male MDPV | 10 | 35.7 (-15.4, 86.7) |
| Intermittent | Female cocaine | 12 | 13.8 (-4.5, 32.0) |
|  | Male cocaine | 10 | 0.3 (-23.3, 23.8) |
|  | Female MDPV | 11 | 17.5 (-86.4, 121.3) |
|  | Male MDPV | 11 | -6.3 (-42.2, 29.7) |
| Short | <1 | 14 | 0.1 (-5.1, 5.3) |
|  | <2 | 18 | 1.2 (-10.6, 13.1) |
|  | 2+ | 5 | -22.9 (-121.0, 75.3) |
| Long | <1 | 10 | **40.2 (4.1, 76.2) \*** |
|  | <2 | 22 | 61.0 (-43, 165.2) |
|  | 2+ | 5 | 164.5 (-84.9, 413.9) |
| Intermittent | <1 | 25 | 0.3 (-16.9, 17.5) |
|  | <2 | 14 | 0.7 (-31.5, 32.8) |
|  | 2 | 5 | 54.8 (-219.8, 329.4) |
| Short |  | 37 | -2.5 (-13.3, 8.4) |
| Long |  | 37 | **69.4 (3.7, 135.1) \*** |
| Intermittent |  | 44 | 6.6 (-18.4, 31.6) |

\* = significant escalation (confidence intervals do not overlap with 0)

## Results

### Acquisition of cocaine and MDPV self-administration

Nearly all rats met acquisition criteria (i.e., ≥ 20 infusions and ≥ 80% of responses on the active lever) within the 14-session acquisition period. Although rats self-administering MDPV acquired earlier (∼4.5 sessions) than rats self-administering cocaine (∼7.5 sessions), the rate of MDPV and cocaine acquisition did not vary by sex (Table 1). The level of acquisition (i.e., mean infusions earned during sessions 12-14) did not differ as a function of either sex or drug (Table 1). Retrospective analyses revealed that rats with higher phenotype scores at the initial phenotyping period acquired more quickly and earned more infusions than rats with lower phenotype scores (Table 1).

### Phenotype 1

After the initial phenotyping period (i.e., ∼5 weeks of short access self-administration), a two-factor (drug x sex) ANOVA revealed main effects of both sex (F [1, 114] = 5.91; p=0.017) and drug (F [1, 114] = 26.09; p<0.0001), where females had higher scores than males, and rats that self-administered MDPV had higher phenotype scores than rats that self-administered cocaine (Figure 1E). This trend was generally true for each of the individual phenotype endpoints where analysis of the mean number of infusions found main effects of sex (F [1, 114] = 7.38; P=0.0076), with females earning more infusions than males, and drug (F [1, 114] = 9.28; P=0.0029), where more infusions of MDPV were self-administered than cocaine (Figure 1A). Similarly, analysis of pre-session TO responses revealed females made more responses than males (F [1, 114] = 10.40; P=0.0016) and rats self-administering MDPV made more responses than rats self-administering cocaine (F [1, 114] = 36.56; P<0.0001) (Figure 1B). Analysis of intercomponent TO responses found that females responded more than males (F [1, 114] = 7.47; P=0.0073) and rats self-administering MDPV responded more than rats self-administering cocaine (F [1, 114] = 42.63; P<0.0001) (Figure 1C). In contrast, though rats self-administering MDPV had higher punishment scores than rats self-administering cocaine (F [1, 114] = 10.28; P=0.0017), there was no main effect of sex (F [1, 114] = 0.22; P=0.6436) (Figure 1D). Importantly, the behavioral response to noncontingent footshock did not differ as a function of SUD-like phenotype score or self-administration drug (phenotype score: F [2, 115] = 0.17); P=0.8453; drug: F [1, 116] = 0.17; P=0.6795) (Supplemental Figure 1), however, females were more sensitive than males (F [1, 116] = 11.27; P=0.0011) (Supplemental Figure 2). There were no sex x drug interactions for any behavioral endpoints or the overall phenotype score.

### Access conditions

The mean number of infusions and rate of responding over the 21-day access condition manipulation are shown in Figure 2. 3-factor ANOVA (access condition x drug x sex) revealed significant main effects of access for both number of infusions earned (F [2, 106] = 108.40; P<0.0001), where long > intermittent > short, and rate of responding (F [2, 106] = 41.14; P<0.0001), where intermittent > long = short. Consistent with the first phenotyping period, rats that self-administered MDPV earned more infusions (F [1, 106] = 13.70; P=0.0003) and responded at a faster rate [F (1, 106] = 13.76; P=0.0003) than rats that self-administered cocaine. Similarly, females earned more infusions (F [1, 106] = 10.23; P=0.0018) and responded at a faster rate (F [1, 106] = 10.22; P=0.0018) than males. Rats that self-administered cocaine, but not MDPV, under long-access conditions showed a significant escalation in drug intake (Table 2).

### Phenotype 2

Redeterminations of the SUD-like phenotype score after manipulating access condition are shown in Figure 3. There were no main effects of access condition on any of the behavioral endpoints nor the overall phenotype score. However, as was observed during the initial phenotyping period, there were main effects of sex and drug on multiple endpoints, as well as the overall phenotype score. Rats that self-administered MDPV earned more infusions (F [1, 106] = 20.58; P<0.0001), made more pre-session TO responses (F [1, 106] = 15.14; P=0.0002), made more intercomponent TO responses (F [1, 106] = 19.22; P<0.0001), and had an overall higher phenotype score than rats self-administering cocaine (F [1, 106] = 11.73; P=0.0009). Females earned more infusions (F [1, 106] = 8.71; P=0.0039), made more pre-session TO responses (F [1, 106] = 5.24; P=0.0240), and had higher phenotype scores (F [1, 106] = 5.36; P=0.0226) compared to males. Though there was no effect of access condition on the overall phenotype score, analysis of the change in phenotype score (Figure 3F) revealed that long-access to cocaine resulted in a significant increase in phenotype score for female rats (mean: 0.44; 95% CI: 0.07-0.82). After self-administration concluded, sensitivity to noncontingent footshock was measured. There was no difference between rats with low, mid, and high phenotype scores or between rats self-administering MDPV or cocaine; however, female rats were more sensitive than male rats (Figure S1).

### Cue reactivity

The number of responses made during the cue reactivity test (i.e., for drug-paired stimuli and a saline infusion) is shown in Figure 4. There were significant main effects of access condition (F [2, 106] = 5.21; P=0.0069), drug (F [1, 106] = 5.05; P=0.0268), and sex (F [1, 106] = 5.74; P=0.0183), where rats that previously self-administered MDPV made more responses than rats that had self-administered cocaine, and females made more responses than males (Figure 4A). Post-hoc analyses revealed that rats with a history of intermittent-access self-administration also made more responses than rats that self-administered under short-(P=0.0126) or long-access (P=0.0290) conditions; responding by short– and long-access rats did not differ (P=0.9538). There were no significant interactions (P≥0.2526). When cue reactivity responses were analyzed by phenotype score, there were no significant main effects (P≥0.0774) or interactions (P≥0.2424) (Figure 4B).

### Measures of receptor sensitivity or availability

Behavioral responses (yawning) to non-contingent administration of pramipexole, a dopamine D_3_/D_2_ receptor agonist, and lorcaserin, a 5-HT_2C_ receptor agonist, were evaluated in a subset of rats (n=43), both prior to initiating self-administration and again after the cue-reactivity tests. Pramipexole dose-dependently induced yawning in male and female rats, although females yawned about half as much as males (Figure S2). There were no effects of phenotype score at either time point; however, there was a main effect of time on the composite yawning score (minimally effect dose x peak number of yawns) in both female (F [1, 63] = 14.70; P=0.0003) and male rats (F (2, 59) = 19.46; P<0.0001) where rats had a higher composite yawning score after self-administration compared to before self-administration began (Figure S2). Lorcaserin did not reliably induce yawning in most rats (data not shown).

Quantitative autoradiography studies were conducted on brain tissue collected from a subset of rats (n=60) after the cue reactivity tests. Expression levels of the dopamine transporter and dopamine D_2_, dopamine D_3_, 5-HT_1B_, 5-HT_2A_, or 5-HT_2C_ receptors did not vary as a function of phenotype score or sex in the nucleus accumbens or caudate putamen (Figures S3, S4, S5). However, there was an effect of access condition (intermittent > short) and drug (MDPV > cocaine) for increased 5-HT_1B_ and 5-HT_2C_ receptor expression, respectively (Figures S4, S5, S6).

## Discussion

Similar to the heterogeneous manifestation of SUD in people, rats can develop different levels of SUD-related behaviors. Studying rats with more extreme phenotypes may provide a more translational framework to understand factors that underlie the transition from regular to disordered patterns of substance use. Though a relatively small subset of rats (17-22%) develop the most severe SUD-like phenotype when they are allowed to self-administer cocaine [3,8], a much larger proportion of rats (∼30-40%) engage in aberrantly high levels of drug-taking when MDPV is available for self-administration [25–29]. Thus, the primary goals of the current studies were to directly compare the SUD-like phenotype in male and female rats self-administering MDPV or cocaine, and to determine how manipulating access condition (short-, long-, and intermittent-access) impacted these SUD-like phenotypes. The first central finding was that rats that self-administer MDPV have a more robust SUD-like phenotype than rats that self-administer cocaine after an initial period of short-access self-administration (Figure 1). Second, female rats exhibit a more robust phenotype than male rats (Figures 1, 3). Third, providing rats with long– or intermittent-access to MDPV or cocaine self-administration did not alter the severity of their SUD-like phenotype, except for female rats self-administering cocaine under long-access conditions, which had increased scores during the second phenotyping period (Figure 3). Finally, evidence from behavioral and quantitative autoradiography studies suggests that these differences may not be due to shifts in expression level of DAT, dopamine D_2_ or D_3_ receptors, or 5-HT_1B_, 5-HT _2A_, or 5-HT _2C_ receptors (but see SI discussion).

Consistent with previous studies reporting unusually high levels of drug-taking in male and female rats that self-administer MDPV, rats that self-administered MDPV had a more severe SUD-like phenotype score than rats that self-administered cocaine, regardless of access condition or duration of self-administration. This was primarily due to the increase in infusions earned, responses made during the pre-session time out, and responses made when drug was signaled to be unavailable, replicating and extending our previous studies with high-responder rats [25–29]. Given that sensitivity to punishment frequently contributes to severe SUD-like phenotype [3,4,6–8] and considering some rats that self-administered MDPV had very high punishment scores, it was somewhat unexpected that rats that self-administered cocaine or MDPV did not differ with regard to the punishment endpoint. This was especially surprising given that the punishment score incorporated both footshock sensitivity (IC_50_) and total current received, and some rats self-administering MDPV earned several dozen more infusions than rats self-administering cocaine. Perhaps the footshock schedule (unsignaled, and unable to be avoided without suppressing all responding) masked any differences between the groups and another procedure (e.g., signaled footshock) would tease apart differences between rats self-administering MDPV or cocaine. Alternatively, the phenotype that leads to sensitivity to footshock punishment may be related less to the other behavioral endpoints [67].

Women initiate drug use later than men, but transition from initial substance use to treatment-seeking in a shorter time period and use similar amounts of cocaine as men [68–71], suggesting women may develop a SUD more rapidly and/or with greater severity compared to men. Even though females and males acquired responding for MDPV and cocaine at similar rates and to similar levels, females self-administering either MDPV or cocaine exhibited more severe SUD-like phenotypes than males during both phenotyping periods. Female rats were more sensitive to noncontingent footshock, suggesting their punishment score and, by extension, overall phenotype score may have even been underestimated.

Decades of work suggest that providing rats long periods of access to cocaine self-administration can result in the development of behaviors thought to more closely resemble SUDs in people (e.g., escalated drug intake, resistance to punishment by footshock) [11–16]. More recently, the intermittent access procedure has been shown to promote rapid, binge-like patterns of cocaine use, and increase the reinforcing effectiveness of cocaine [6,17–20]. Both phenomena were observed in the present study, although the escalation was not statistically significant in rats that self-administered MDPV under long-access conditions. Unexpectedly, we found that some rats self-administered up to 80 infusions of MDPV in a single 5-min period during the intermittent-access procedure. Five of 22 of the rats (23%) that self-administered MDPV earned an average of 17-45 infusions per 5-min period across the entire 21-session access manipulation, which is much higher than the approximate 3-12 cocaine infusions in a 5-min period that we and others have observed when unit-doses ranging from 0.25 to 0.4 mg/kg/infusion are available [72–76]. This finding strongly supports the notion that binge-like patterns of dysregulated drug-taking develop in a subset of rats that self-administer MDPV, consistent with what has been reported by humans using MDPV and related synthetic cathinones. Though the access manipulations produced robust behavioral differences, these effects did not carry over into the phenotyping period, suggesting that the changes in patterns of drug-taking induced by long– and/or intermittent-access may not be long-lasting and may be more a function of the schedule of reinforcement than a fundamental change in the “state” of the rat. Though we have previously reported that ‘high-responder” rats will earn significantly more infusions of MDPV under both FR and progressive ratio schedules of reinforcement, we did not evaluate responding under a progressive ratio schedule of reinforcement or use behavioral economics. Other studies find that rats with a history of long– or intermittent-access find cocaine and other reinforcers more reinforcing than rats with a history of short-access self-administration [14,18,72,75,77–82], but see [83].

Though rats with a history of self-administering cocaine under long-compared to short-access conditions have been reported to make more responses during reinstatement or cue reactivity tests [14–16], this effect was not seen in the present study. However, rats that self-administered under intermittent-access did make more responses compared to the other access conditions, consistent with other reports [75,76,84]. We also found that female rats made more responses than male rats during the cue reactivity test, which may be related to the higher rates of relapse and drug craving in women compared to men [85–88]; however, many studies do not report sex differences in reinstatement or cue reactivity tests [14,89,90], but see [91]. These differences could be due to procedural differences (e.g., extinction sessions or a history of punishment).

The differences in MDPV and cocaine at the first phenotyping period could have represented a quicker transition to a SUD-like phenotype; however, the fact that the phenotype scores for cocaine and MDPV did not converge after access suggests that there is something fundamentally different about the development of SUD-like phenotypes in response to MDPV and cocaine self-administration. Since we did not observe any consistent effects of access condition, this study cannot rule out the involvement of DAT or any of the receptors quantified here in the presentation of SUD-like phenotype(s). Additionally, because all rats showed a leftward shift in the pramipexole-induced yawning dose-response function (Figure S2), the assay may not have been sensitive enough to detect relatively small differences in the size of the shift. Future studies could use alternative approaches, such as RNA sequencing or genome-wide association studies [92–95] to cast a wider net to identify underlying factors that contribute to the individual differences in SUD-like phenotype.

The within-subject design can be powerful to evaluate individual differences, with some caveats. For instance, more sessions were spent self-administering under short-access conditions for all groups than the access condition manipulation (70 short-access vs 21 of long– or intermittent-access), which may have attenuated the effects of the access condition. However, we calculated the second phenotype score using data from only the first three sessions following the access condition manipulation and did not find any differences compared to using the average of the entire phenotyping period (data not shown). Additionally, the effects of intermittent-access may have been underestimated since two of the phenotype endpoints (pre-session responses and intercomponent timeout responses) measured responding during signaled periods of unavailability, and rats with intermittent-access had extended periods (5-hours/session) of drug unavailability, which rats in the short– and long-access groups did not experience. However, there was also no effect of intermittent-access on the other two endpoints (i.e., infusions, punishment score), suggesting an overall lack of effect of intermittent-access.

Synthetic cathinones have been reported to produce stimulant and euphoric effects in humans [96], and in the current study, even relatively brief periods of short-access to MDPV self-administration produced high levels of drug-taking and –seeking. Though the SUD-like phenotype established by MDPV was not exacerbated by a history of long– or intermittent-access to MDPV self-administration, it is equally interesting and important to note that even long– or intermittent-access to cocaine was unable to produce an SUD-like phenotype comparable to that established with MDPV. Exploiting the severe phenotype developed in rats self-administering MDPV to investigate the mechanisms that underly the development of the phenotype can provide valuable insight into the transition in people from recreational use to SUD and help identify novel pharmacotherapies for SUD treatment.

## Funding

This work was supported by the National Institutes of Health, including National Institute of Drug Abuse [R01 DA039146 (GTC), R36 DA050955 (MRD), R21 DA046044 (LCD), and R01 DA055703 (LCD)], the jointly-sponsored National Institutes of Health Predoctoral Training Program in the Neurosciences [Grant T32 NS082145 (MRD)], and the Intramural Research Programs of the National Institute on Drug Abuse and National Institute of Alcohol Abuse and Alcoholism [Z1A-DA000527 (KCR)]. It was also supported by the John L Santikos Charitable Foundation endowment to the San Antonio Area Foundation (GGG).

## Competing Interests

The authors have nothing to disclose.

## Acknowledgments

The authors would like to thank Drs. Amy Hauck Newman and Jianjing Cao for providing VK4-116, which was used in some of the quantitative autoradiography studies.

## Author contributions

MRD and GTC conceived the project and designed the experiments with input from GGG and LCD. MRD, MSAB, MS, VA, and MD performed the behavioral experiments. NMB analyzed and scored the yawning videos and contributed to data analysis. MRD, GGG, MSAB, and MS performed quantitative autoradiography experiments and analysis. MRD analyzed and interpreted the experimental data. GTC, KMS, GGG and LCD supervised the research. KCR contributed reagents. MRD and GTC wrote the manuscript with contributions from all authors. All authors read and approved the final manuscript.

## Figure Legends

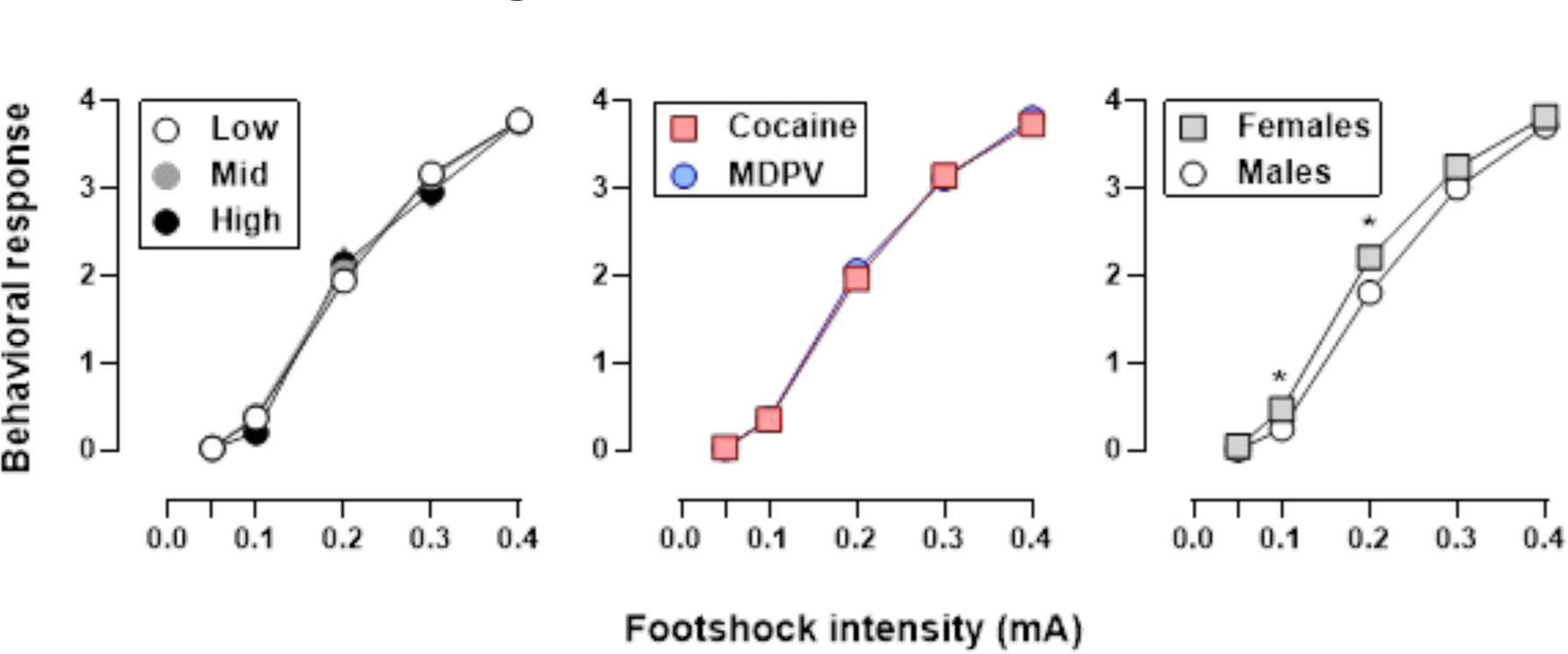
Supplemental Figure 1

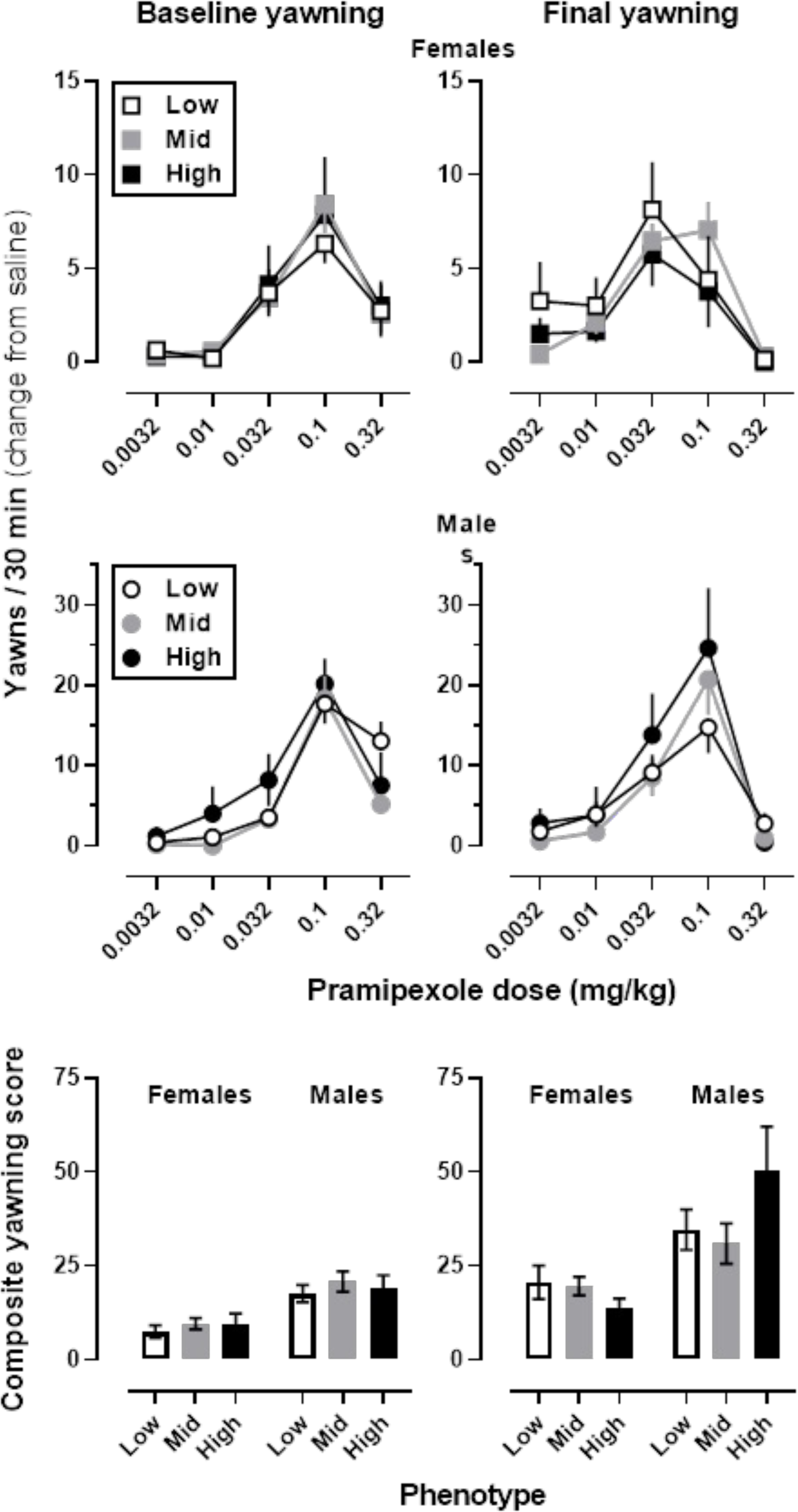
Supplemental Figure 2

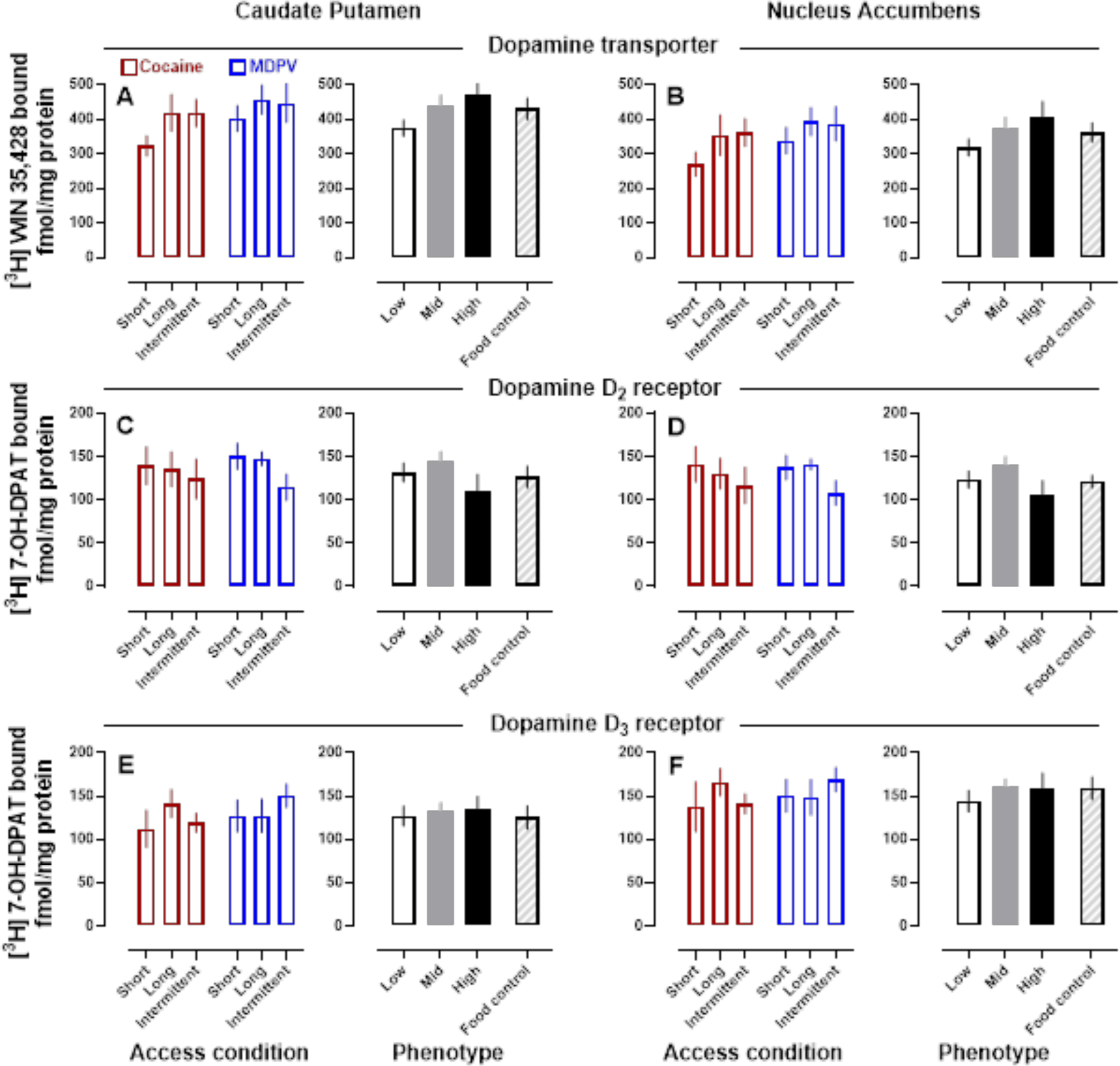
Supplemental Figure 3

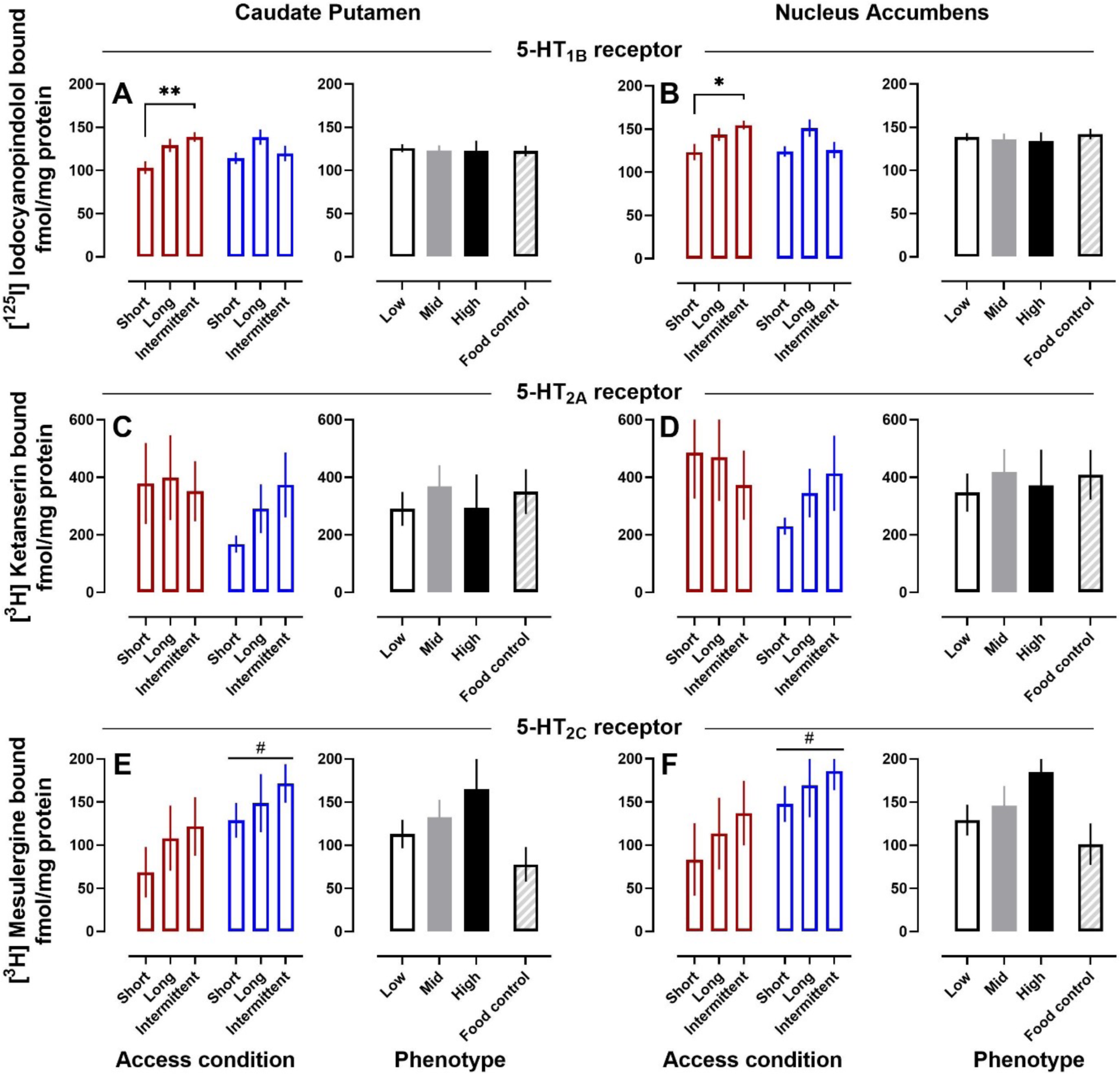
Supplemental Figure 4

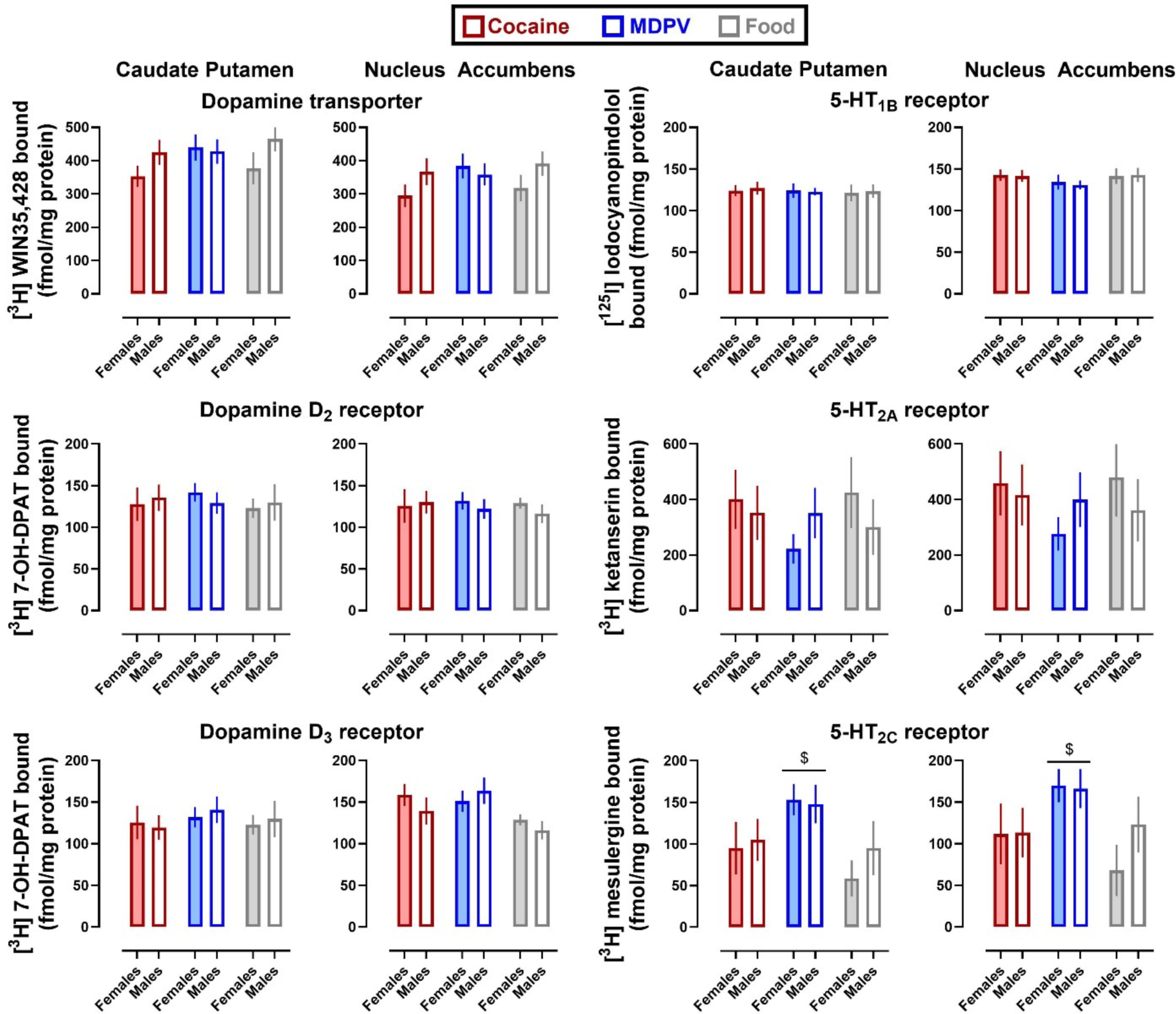
Supplemental Figure 5

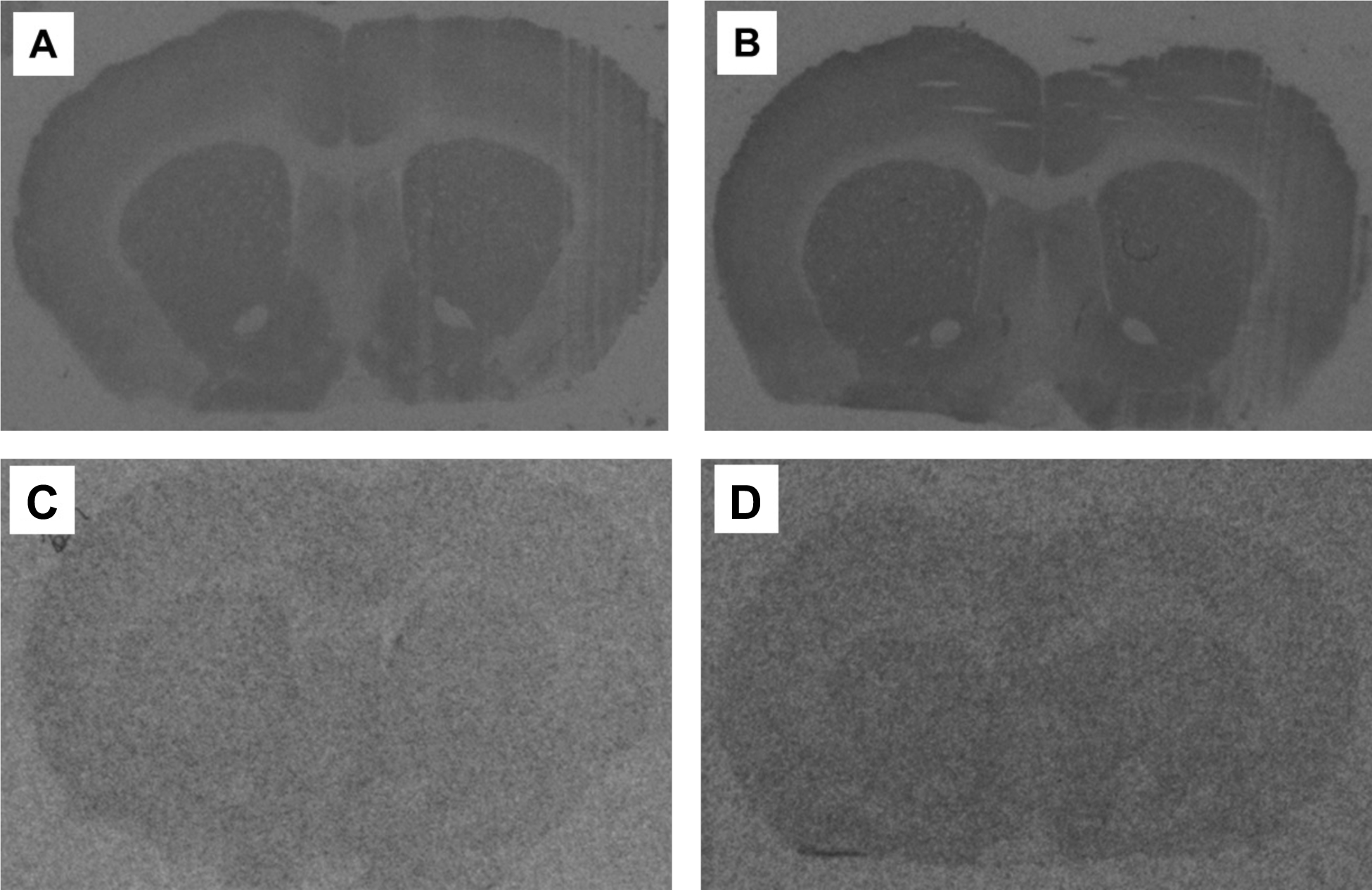
Supplemental Figure 6

## Notes

### Competing Interest Statement

The authors have declared no competing interest.

