## Supplemental Materials for "Effects of access condition on substance use disorder-like phenotypes in male and female rats self-administering MDPV or cocaine"

### Supplemental Methods

#### *Behavioral Observations of Yawning*

A yawn was defined as a broad, prolonged (~1 sec) opening, then rapid closure of the mouth [1-3]. Prior to observations, rats were weighed and then transferred to clear plastic enclosures (Model 435; IITC Inc. Life Science, Woodland Hills, CA) on a wire mesh grid to habituate for 30-60-min before injections. All rats were first habituated to the experimental procedure and received 6 injections of saline (1 ml/kg; intraperitoneally [i.p.]) at least one day before the first experimental observations of yawning. Rats were pseudo-randomly split into two groups, where a subset of rats only ever received injections of saline (n=45, for quantitative autoradiography experiments) and the remaining rats received pramipexole and lorcaserin injections (n=73), with the order of drugs counterbalanced across rats. Yawning observations always occurred in pairs (e.g., two observations before self-administration and two observations after the drug-free period); rats were counterbalanced for whether pramipexole or lorcaserin were administered during the first or second observation period, but the order was consistent across testing periods for each rat.

On observation days, the first injection for all rats was saline (1 ml/kg; i.p.), followed by 5 additional injections (i.p.) of saline, cumulative doses of pramipexole (0.0032-0.32 mg/kg) or cumulative doses of lorcaserin (0.01-1 mg/kg) every 30-min [1-4]. A mirror was placed behind the plastic enclosures to allow for video recordings of behavior, which were scored using Simple Video Coder [5] by blinded observers (NMB, VA and MD) at a later time. Yawns during each of the 30-min periods after injections were counted, and all yawning experiments were conducted between 12:00 and 19:00.

#### *Quantitative Autoradiography*

##### *Drug-Free Control Rats*

A subset of rats (n=16) went through the entire experiment as described, but responded for 1 grain-based pellet (Dustless Precision Pellets® Rodent, 45-mg; Bio-Serv, Flemington, NJ) instead of responding for cocaine or MDPV to serve as drug-free controls with operant history for the quantitative autoradiography portion of these studies. Half of the rats (n=4 males; n=4 females) responded under short-access conditions; the remaining rats were assigned to long- (n=2 males; n=2 females) or intermittent-access (n=2 males; n=2 females).

##### *Preparation*

Rats were euthanized by rapid decapitation without anesthesia. Brains were extracted and immediately frozen on dry ice before being stored in a -80°C freezer until sectioning. Brains were sectioned using a cryostat and 20 μm coronal sections were collected to examine the caudate putamen and nucleus accumbens due to the role of these brain regions in reinforcement. Sections were thaw-mounted on gelatin-coated microscope slides, then desiccated at 4°C overnight before returning to the -80°C freezer for storage until all tissue was sectioned before beginning quantitative autoradiography studies. Slides were thawed for 20-min at 4°C before incubation steps, detailed below.

*Dopamine Transporter Autoradiography Using [$^3$H] WIN 35,428*

Slides were pre-incubated in 30 mM sodium phosphate buffer (pH 7.4) at room temperature for 1-hour before a 2-hour incubation at 4°C in the same buffer but in the presence of 10 nM (approximately 5 times the dissociation constant [$K_D$]) [N-methyl-$^3$H] WIN 35,428 (specific activity: 82.5 Ci/mmol; PerkinElmer NET1033250UC, Lot: 2795346, Boston, MA). Non-specific binding was quantified in the presence of 10µM GBR12909. Slides were then washed twice (1-min each) in ice cold 30 mM sodium phosphate (pH 7.4), followed by a 1-sec dip in ice cold deionized (di) $H_2O$. Methods modified from [6,7].

*Dopamine $D_2$ and $D_3$ Receptor Autoradiography Using [$^3$H]7-OH-DPAT*

Slides were pre-incubated in 50 mM Tris hydrochloride, 1 mM EDTA, 5 mM magnesium chloride buffer (pH 7.4) at room temperature for 30-min before a 1.5-hour incubation at 4°C in the same buffer but also in the presence of 10 nM [$^3$H] 7-hydroxy-DPAT (~3 times the $K_D$; 100 Ci/mmol; PerkinElmer NET1169250UC, Lot: 2716021, Boston, MA). For dopamine $D_2$ receptor autoradiography, the buffer also contained 1 µM VK4-116 to prevent ligand binding to dopamine $D_3$ receptors. For dopamine $D_3$ receptor autoradiography, the buffer also contained 10 µm L-741,626 to block ligand binding to dopamine $D_2$ receptors. In both cases, non-specific binding was quantified in the presence of 10 µM haloperidol. Slides were then washed thrice (5-min each) in ice cold 50 mM Tris buffer (pH 7.4), followed by a 1-sec dip in ice cold di$H_2O$. Methods modified from [8,9].

*5-$HT_{2A}$ Receptor Autoradiography Using [$^3$H] Ketanserin*

Slides were pre-incubated in 170 mM Tris buffer (pH 7.5) at room temperature for 15-min before incubating for 1-hour at room temperature in the same buffer but in the presence of 20 nM (5 times the $K_D$) [ethylene-$^3$H] ketanserin hydrochloride (47.3 Ci/mmol; PerkinElmer NET791250UC, Lot: 2233253, Boston, MA) and 1 µM prazosin (to prevent ligand binding to α1 adrenergic receptors). Non-specific binding was quantified in the presence of 10 µM M100,907. Slides were then washed twice (10-min each) in ice cold 170 mM Tris (pH 7.6), followed by a 1-sec dip in ice cold di$H_2O$. Methods modified from [10].

*5-$HT_{2C}$ Receptor Autoradiography Using [$^3$H] Mesulergine*

Slides were pre-incubated in 170 mM Tris buffer (pH 7.5) at room temperature for 40-min before a 2-hour incubation at room temperature in the same buffer but in the presence of 5 nM (5 times the $K_D$) [N-methyl-$^3$H] mesulergine (76.8 Ci/mmol; PerkinElmer NET114825UC, Lot: 2834417, Boston, MA) and 100 nM M100,907 (to prevent binding of ligand to 5-$HT_{2A}$ receptors). Non-specific binding was quantified in the presence of 1 µM RS 102221 hydrochloride. Slides were then washed twice (10-min each) in ice cold 170mM Tris buffer (pH 7.6), followed by a 1-sec dip in ice cold di$H_2O$. Methods modified from [10].

*5-HT$_{1B}$ Receptor Autoradiography Using [$^{125}$I] Iodo-(±)-cyanopindolol*

Slides were pre-incubated in 170 mM Tris buffer (pH 7.4) at room temperature for 15-min before a 2-hour incubation in the same buffer, but in the presence of 0.01% ascorbic acid, 10 µM (-) isoproterenol to prevent radioligand binding to β1 adrenergic binding, and 30 nM [$^{125}$I] iodo-(±)-cyanopindolol (5 times the $K_D$; PerkinElmer, NEX174100UC, Lot COA0510, Boston, MA). Non-specific binding was quantified in the presence of 1 µM GR127935. Slides were washed twice (10-min each) in ice cold 170 mM Tris buffer, followed by a 1-sec dip in ice cold di$H_2O$. Methods modified from [11].

*Imaging*

After being rinsed in di$H_2O$, slides were dried and exposed to Kodak Biomax MR film for 1 day (5-HT$_{1B}$), ~6 weeks (5-HT$_{2A}$), ~8 weeks (DAT, 5-HT$_{2C}$, dopamine D$_2$), or ~16 weeks (dopamine D$_3$) with [$^3$H] standards (1 x 0.02 mCi/slide; ART-123, American Radiolabeled Chemicals, Inc., St Louis, MO). Autoradiographic images were captured on a digital imaging station (Northern Lights illuminator, Scion Image CCD camera, copy stand) and calibrated using a linear function. Binding density was measured using Image J software (http://rsb.info.nih.gov/ij/download.html) by one or two, trained, blinded observers (MSAB and MS; inter-observer variability: $R^2 > 0.99$).

### ***Drugs***

Pramipexole was purchased from Sigma Aldrich (St. Louis, MO) and lorcaserin was purchased from MedChem Express (Monmouth Junction, NJ). Both were dissolved in sterile saline and administered i.p. at a volume of 1 ml/kg.

L-741,626, M100,907, prazosin hydrochloride, (-) isoproterenol, and GBR12909 were purchased from Sigma Aldrich (St. Louis, MO). RS 102221 hydrochloride was purchased from Tocris (Ballwin, MO) and GR127935 was purchased from Calbiochem (San Diego, CA). VK4-116 was synthesized and supplied by Jianjing Cao and Amy Hauck Newman. All ligands used in the autoradiography studies were dissolved in dimethyl sulfoxide. The radioligands ([N-methyl-$^3$H] WIN 35,428, [$^3$H] 7-hydroxy-DPAT, [ethylene-$^3$H] ketanserin hydrochloride, [N-methyl-$^3$H] mesulergine, [$^{125}$I] Iodo-(±)-cyanopindolol) were purchased from PerkinElmer (Boston, MA).

### ***Statistical Analyses***

*Yawning Analyses*

The pramipexole-induced yawning data were transformed by subtracting the number of yawns after the saline injection from all other data points within individual subjects. To quantify dopamine D$_3$ receptor sensitivity, a composite yawning score was calculated by multiplying the peak number of yawns by the -log of the first effective dose of yawning. Effects of phenotype score were evaluated by a two-factor (pramipexole dose x phenotype) ANOVA. A 3-factor (time x phenotype score x sex) ANOVA on the composite yawning score was used to evaluate the shift in the dose-response curves overtime

(Supplemental Figure). Lorcaserin-induced yawning data were excluded from analyses as lorcaserin did not reliably induce yawning in a majority of subjects.

*Quantitative Autoradiography Analyses*

The mean nonspecific binding measure was subtracted from the total binding measure for each brain region to obtain specific binding measures. Three-factor (drug x sex x access condition or drug x sex x phenotype score) ANOVAs were performed on the measures of specific binding in Figures S3 and S4. Two-factor (reinforcer x sex) ANOVAs were performed in Figure S5. Tukey's post-hoc analyses were performed when there was a significant main effect of access condition.

### Supplemental Results

#### *Quantitative Autoradiography*

DAT binding in the caudate putamen (Figure S3A) and nucleus accumbens (Figure S3B) was measured. There were no significant effects of drug, access condition, or phenotype score. There was no measurable amount of nonspecific binding for DAT. Dopamine $D_2$ receptor binding in the nucleus accumbens (Figure S3C) and caudate putamen (Figure S3D) was measured. There were no significant effects of drug, access condition, or phenotype score. There was no measurable amount of nonspecific binding for dopamine $D_2$ receptors. Dopamine $D_3$ receptor binding in the caudate putamen (Figure S3E) and nucleus accumbens (Figure S3F) was measured. There were no significant effects of drug, access condition, or phenotype score. There was no measurable amount of nonspecific binding for dopamine $D_3$ receptors. There were no effects of sex for any of the dopaminergic targets (Figure S5).

In contrast, in caudate putamen (Figure S4A) and nucleus accumbens (Figure S4B) there were significant main effects of access condition (caudate putamen: $F [2, 39] = 5.699$; $P=0.0067$; nucleus accumbens: $F [2, 39] = 4.275$, $P=0.0210$) on 5-$HT_{1B}$ receptor binding. Post-hoc analyses revealed that rats self-administering under cocaine short-access conditions had significantly lower 5-$HT_{1B}$ receptor binding in the caudate putamen than rats that self-administered cocaine under intermittent-access conditions in the caudate putamen ($P=0.0094$) and nucleus accumbens ($P=0.036$). Nonspecific binding was measured at 29.2 ± 9.5 fmol per mg of protein. Nonspecific binding was measured at 29.2 ± 9.5 and 29.1 ± 9.2 fmol per mg of protein in the caudate putamen and nucleus accumbens, respectively. Radiograms for short- and intermittent-access are shown in Figure S6.

5-$HT_{2A}$ receptor binding was measured in the caudate putamen (Figure S4C) and nucleus accumbens (Figure S4D). There were no significant effects of drug, access condition, or phenotype score. Nonspecific binding for [$^3$H] ketanserin was measured at 991.0 ± 198.5 fmol per mg of protein and 775.9 ± 179.7 fmol per mg of protein in the caudate putamen and nucleus accumbens, respectively. 5-$HT_{2C}$ receptor binding was measured in caudate putamen (Figure S4E) and nucleus accumbens (Figure S4F), and though there were no significant effects of access condition or phenotype score, there were main effects of drug history. Rats with a history of MDPV self-administration had greater 5-$HT_{2C}$ receptor binding than rats with a history of cocaine self-administration in both the nucleus accumbens ($F [1, 37] = 4.42$; $P=0.0424$) and caudate putamen ($F [1, 37] = 4.42$; $P=0.0425$). Additionally, there was a main effect of reinforcer (caudate putamen: $F [2, 52] = 4.57$; $P=0.0148$; nucleus accumbens: $F [2, 52] = 3.67$; $P=0.0322$), where rats that self-administered MDPV had greater 5-$HT_{2C}$ receptor binding than rats that responded for food in both the caudate putamen ($P=0.0185$) and nucleus accumbens ($P=0.0445$) (Figure S5). Nonspecific binding for [$^3$H] mesulergine was measured at 488.0 ± 85.9 and 504.9 ± 84. 2 fmol per mg of protein in the caudate putamen and nucleus accumbens, respectively. Representative autoradiograms are shown in Figure S6. There was no effect of sex for any of the serotonergic targets (Figure S5)

**Supplemental Discussion**

Dopamine $D_2$ and $D_3$ receptor sensitivity did not predict the SUD-like phenotype (Figure S2), nor did we observe consistent changes in expression of DAT, dopamine $D_2$, dopamine $D_3$ (Figure S3) 5-$HT_{1B}$, 5-$HT_{2A}$, or 5-$HT_{2C}$ receptors (Figure S4) that could be associated with SUD-like phenotype score. Although multiple imaging studies in people and nonhuman primates show consistent downregulation of dopamine $D_2$ receptors after stimulant use [12-17], the effects in rodents are less consistent. For instance, some studies show transient decreases in dopamine $D_2$ receptor expression [18] others show decreases that emerge 45 days, but not 1 day, after the last cocaine administration [19] while others find no change in striatal dopamine $D_2$ receptor expression after cocaine administration [20]. Still other studies suggest there is a downregulation of dopamine $D_2$ receptors after noncontingent/yoked, but not self-administered, cocaine [21]. Though the present findings do not align with the relatively consistent human and nonhuman primate literature, the rodent literature appears to be much more variable and subject to changes in experimental protocol (e.g., route of administration, dose, time period, contingencies).

Most rodent studies that examine changes in dopamine $D_3$ receptor expression do so after a relatively short history of cocaine (e.g., 5-14 days), resulting in much lower cocaine intake (e.g., 45-140 mg/kg; although in one study, total intake was ~600 mg/kg) [2,19,22-24]. In contrast, the current study evaluated changes in dopamine $D_3$ receptor expression after >80 self-administration sessions and ~1300 mg/kg of cocaine intake on average (range: 500-2600 mg/kg). There are mixed reports about the time until increases in dopamine $D_3$ receptor expression are observed, with some studies finding increases 1-3 days after the last cocaine administration [22] while others did not observe increases 1-3 days after, but instead saw increases weeks after last cocaine administration [2,19,23,24]. Because of these results and prior evidence of increases in dopamine $D_3$ receptors in people who use stimulants, it was surprising that we did not observe significant increases in dopamine $D_3$ receptor expression. Perhaps the differences are related to the relatively long drug history and/or high level of drug intake not typically used by other laboratories.

In contrast to dopamine $D_3$ receptors where changes often grow overtime, DAT expression increases after cocaine administration but then appears to normalize after weeks [12,15,25-29]. The discrepancy between our findings and previous results suggests it is likely that we did not observe an increase in DAT expression due to the extended drug-free period used in this study compared to previous work looking at changes within 24 hours of last cocaine administration.

Serotonergic modulation of SUD-related phenotypes has been less thoroughly explored compared to dopaminergic modulation. Overexpression of 5-$HT_{1B}$ receptors results in an increase in drug-taking across a range of cocaine doses [30], and yet knockdown of these receptors results in a more compulsive, or punishment-resistant phenotype [31]. Further research is needed to better understand the changes in receptor expression that may occur as a function of stimulant use, especially given the increase in 5-$HT_{1B}$ receptor expression in rats that self-administered cocaine under intermittent-access conditions compared to short-access in the present studies. It was previously reported that 24-hours after cocaine administration, there is an increase in 5-$HT_{2A}$ receptor expression [32]; however, we found no change in 5-$HT_{2A}$ receptor expression after a 3-week drug free period. This suggests further research is needed to determine the time course of changes in 5-$HT_{2A}$ receptors. Although we are not aware of any reported change in 5-$HT_{2C}$ receptor expression after cocaine self-administration, lower levels of 5-$HT_{2C}$ receptors are associated with increased measures of impulsivity, which, in turn, are associated with greater cocaine self-administration [33]. At the time point we examined, we found a non-significant trend for an increase in 5-$HT_{2C}$ receptor expression as a function of phenotype score. 5-$HT_{2C}$ receptor expression was increased in rats that self-administered MDPV relative to those that responded for cocaine or food; however, the mechanisms underlying these changes are unclear.

Although we found limited or no change in DAT and receptor expression in these studies, we only examined $B_{max}$.; saturation curves might have revealed differences in $K_D$ as a function of phenotype. Future studies could apply other methods including biotinylation assays to examine differences in localization (e.g., intracellular, plasma membrane), and GTP-γ-S binding to get a better idea of functional changes. Though these targets were selected because they have previously been shown to change after a history of stimulant use and/or modulate drug-taking, the receptors and transporter quantified are only a small subset of potential proteins that could mediate difference between rats with a mild and severe SUD-like phenotype.

**Supplemental Figures**

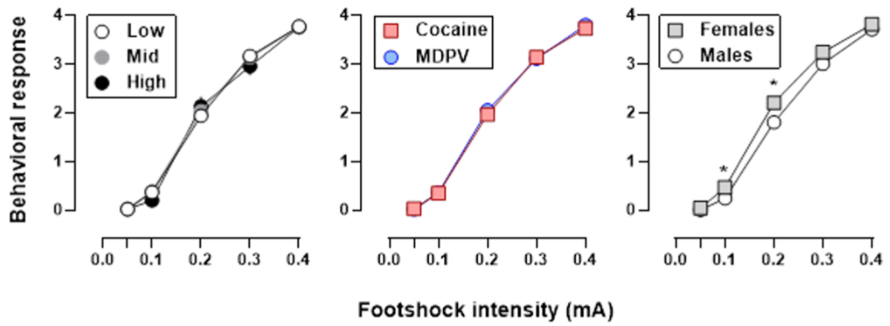


Supplemental Figure 1. Behavioral Response to Noncontingent Shock

Response to non-contingent shock in rats split by phenotype score (left), self-administered drug (middle), or sex (right). Ordinate: behavioral response to noncontingent shock (see methods for details). Abscissa: footshock intensity (mA). Error bars represent ± S.E.M. * P<0.05 different from males

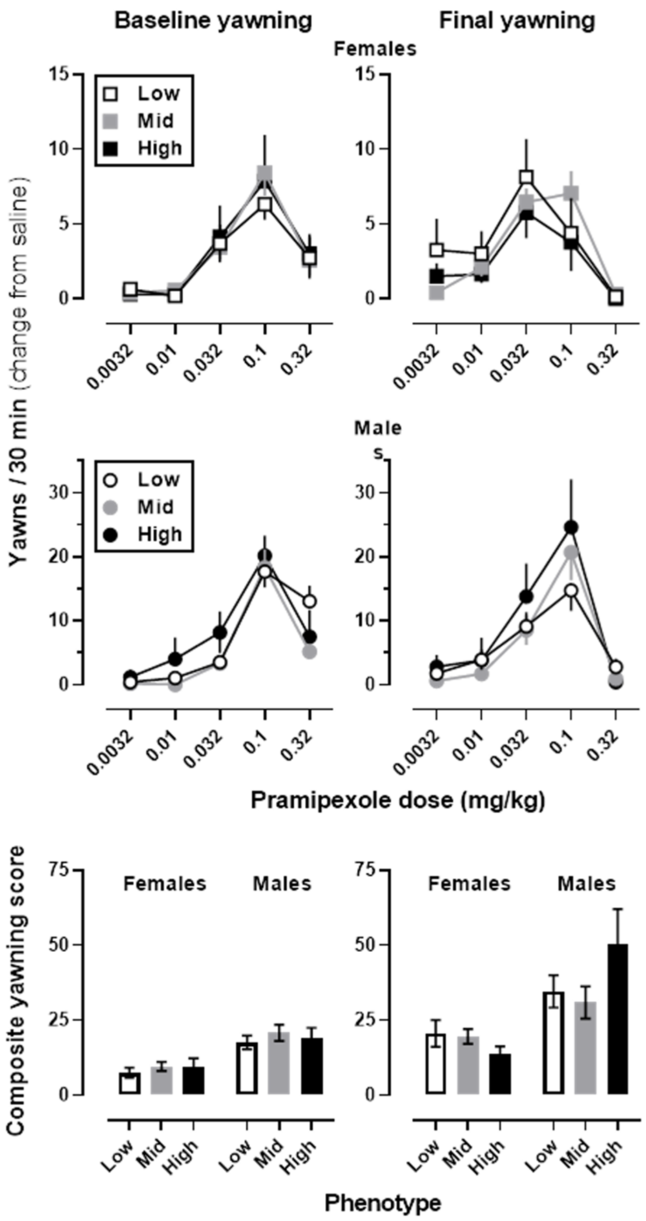


Supplemental Figure 2. Pramipexole-Induced Yawning as a Behavioral Measure of Dopamine $D_3$ Receptor Availability

Yawning dose-response curves generated prior to initiation of self-administration (left) or after the 21-day drug free period (right) in female (squares; top row) and male rats (circles; middle row). Data are separated by low (open symbols), mid (grey symbols), or high phenotype score (black symbols). Ordinate: number of yawns minus number of yawns after saline injection. Abscissa: cumulative dose of pramipexole, expressed in log units. Bottom figure: composite yawning score (see methods for details) by sex, phenotype score and time. Error bars represent ± S.E.M.

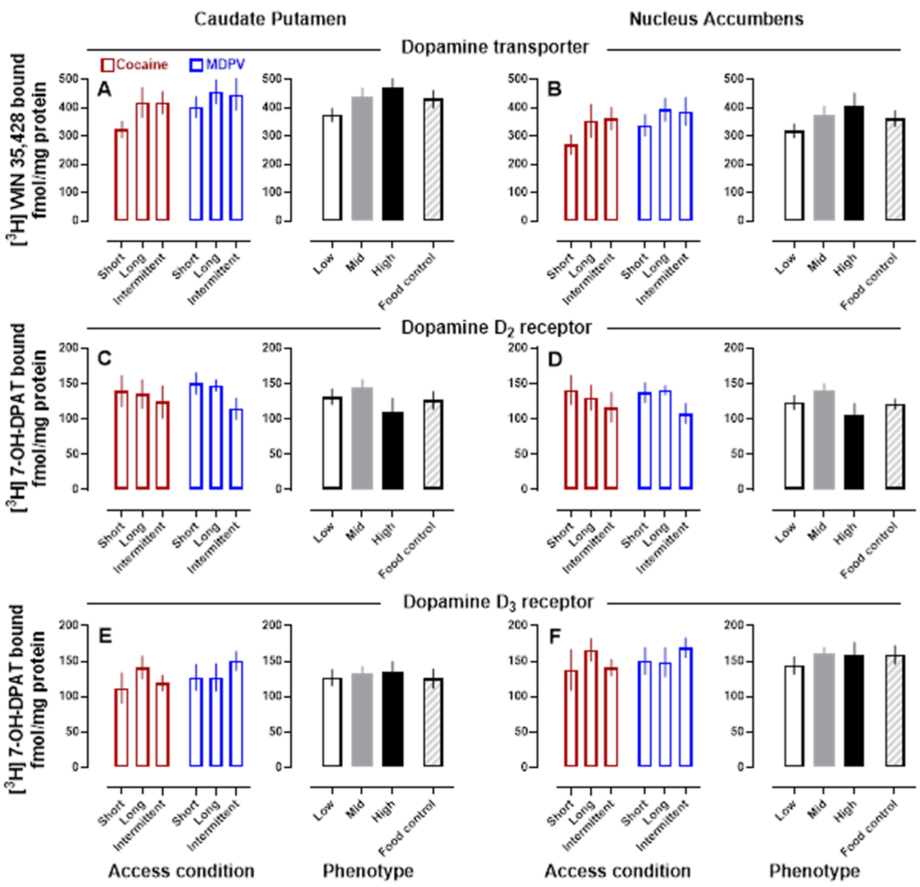


Supplemental Figure 3. Quantitative Autoradiography of Dopaminergic Targets

Dopamine transporter (top row), dopamine $D_2$ (middle row), and dopamine $D_3$ (bottom row) expression in the caudate putamen (left columns) and nucleus accumbens (right columns). Data are separated by drug (cocaine: red; MDPV: blue) and access condition on the left, or SUD-like phenotype score (low: open; mid: grey; high: black) and drug-free food controls (striped) on the right. Error bars represent ± S.E.M.

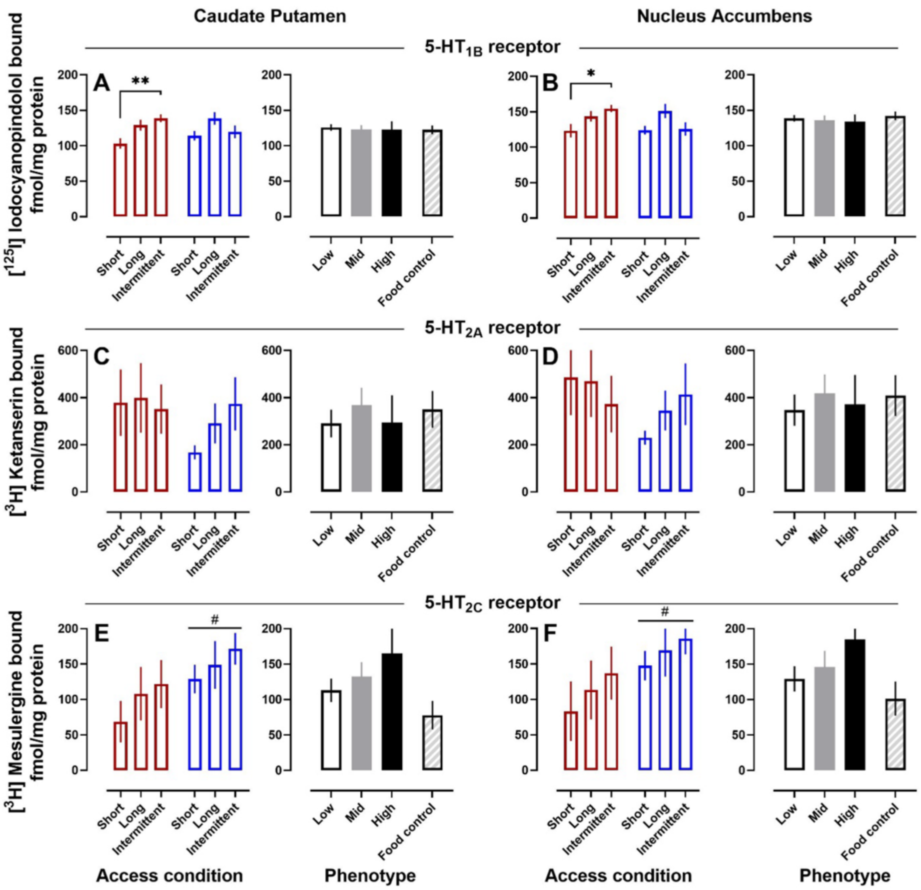


Supplemental Figure 4. Quantitative Autoradiography as a Function of Reinforcer and Sex

Dopamine transporter expression (top left) and dopamine $D_2$ (middle left), dopamine $D_3$ (bottom left), 5-$HT_{1B}$ (top right), 5-$HT_{2A}$ (middle right), and 5-$HT_{2C}$ (bottom right) receptor expression in the caudate putamen (left columns) and nucleus accumbens (right columns). Data are separated by reinforcer (cocaine: red; MDPV: blue; food: gray) and sex (females: filled bars; males: open bars). Error bars represent ± S.E.M. \$=P<0.05 compared to food controls

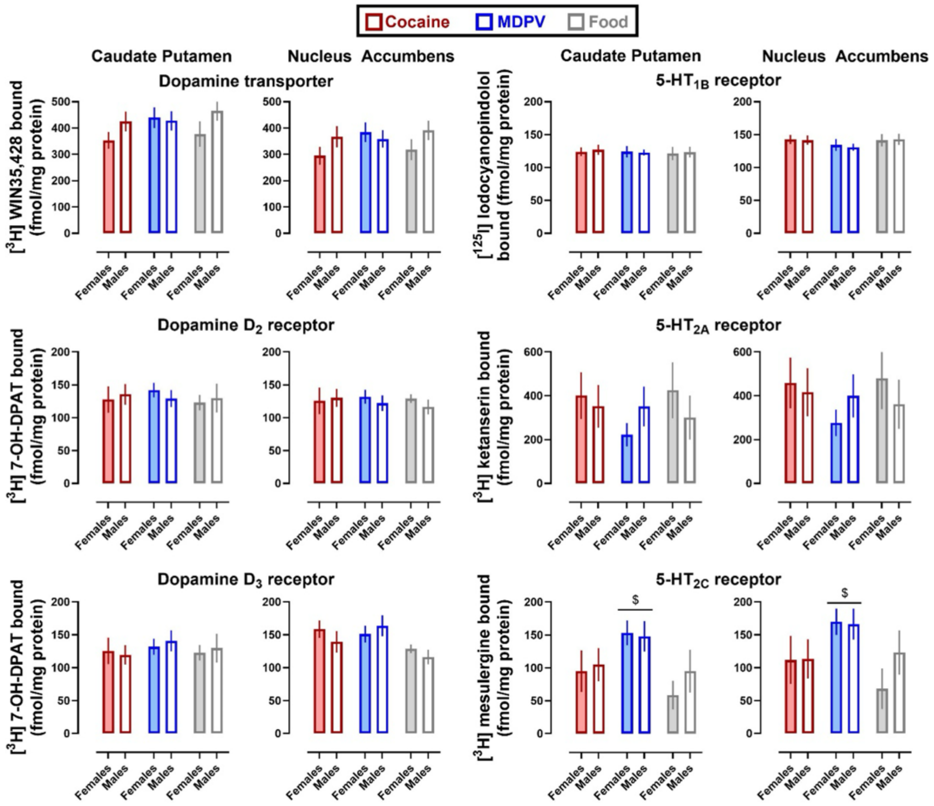


Supplemental Figure 5. Quantitative Autoradiography of Serotonergic Targets

5-$HT_{1B}$ (top row), 5-$HT_{2A}$ (middle row), and 5-$HT_{2C}$ (bottom row) receptor expression in the caudate putamen (left columns) and nucleus accumbens (right columns). Data are separated by drug (cocaine: red; MDPV: blue) and access condition on the left, or SUD-like phenotype score (low: open; mid: grey; high: black) and drug-free food controls (striped) on the right. Error bars represent ± S.E.M *=P<0.05, **=P<0.01, #=main effect of drug where P<0.05

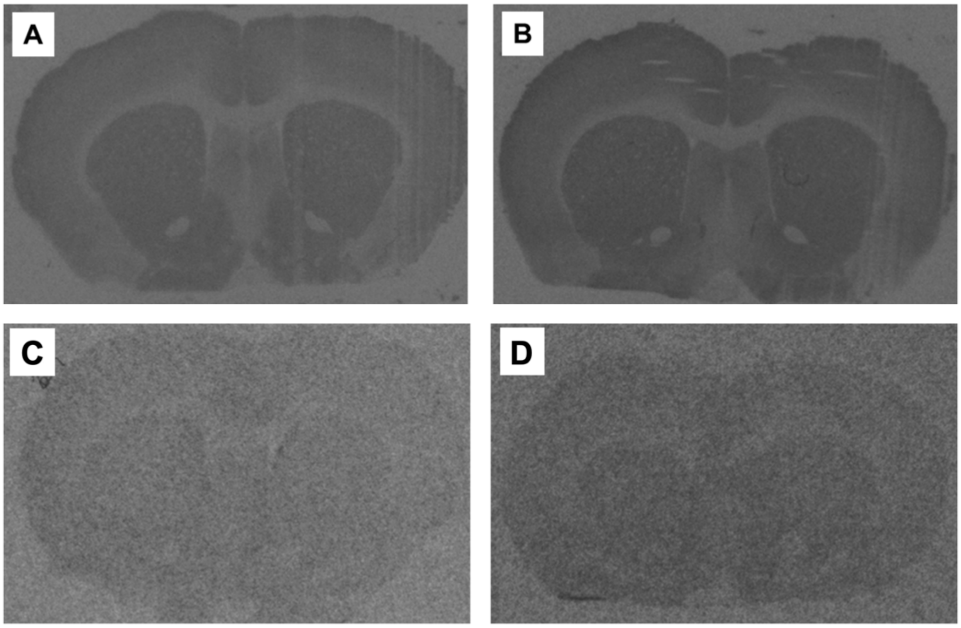


Supplemental Figure 6. 5-$HT_{1B}$ and 5-$HT_{2C}$ Receptor Autoradiograms

Representative autoradiograms of [$^{125}$I] Iodo-(±)-cyanopindolol (30 nM) binding to 5-$HT_{1B}$ receptors in the caudate putamen and nucleus accumbens in rats that self-administered cocaine under short- (A) or intermittent-access (B) conditions. Representative autoradiograms of [$^{3}$H] mesulergine (5 nM) binding to 5-$HT_{2C}$ receptors in the caudate putamen and nucleus accumbens in rats that self-administered cocaine (C) or MDPV (D). Shown are autoradiograms of total binding.
